# Highly dynamic evolution of the chemosensory system driven by gene gain and loss across subterranean beetles

**DOI:** 10.1101/2022.12.08.519422

**Authors:** Pau Balart-García, Tessa M. Bradford, Perry G. Beasley-Hall, Slavko Polak, Steven J. B. Cooper, Rosa Fernández

## Abstract

Chemical cues in subterranean habitats differ highly from those on the surface due to the contrasting environmental conditions, such as absolute darkness, high humidity or food scarcity. Subterranean animals underwent changes to their sensory systems to facilitate the perception of essential stimuli for underground lifestyles. Despite representing unique systems to understand biological adaptation, the genomic basis of chemosensation across cave-dwelling species remains unexplored from a macroevolutionary perspective. Here, we explore the evolution of chemoreception in three beetle tribes that underwent at least six independent transitions to the underground, through a phylogenomics spyglass. Our findings suggest that the chemosensory gene repertoire varies dramatically between species. Overall, no parallel changes in the net rate of evolution of chemosensory gene families were detected prior, during, or after the habitat shift among subterranean lineages. Contrarily, we found evidence of lineage-specific changes within surface and subterranean lineages. However, our results reveal key duplications and losses shared between some of the lineages transitioning to the underground, including the loss of sugar receptors and gene duplications of the highly conserved ionotropic receptors IR25a and IR8a, involved in thermal and humidity sensing among other olfactory roles in insects. These duplications were detected both in independent subterranean lineages and their surface relatives, suggesting parallel evolution of these genes across lineages giving rise to cave-dwelling species. Overall, our results shed light on the genomic basis of chemoreception in subterranean beetles and contribute to our understanding of the genomic underpinnings of adaptation to the subterranean lifestyle at a macroevolutionary scale.

## 1. INTRODUCTION

Chemoreception is an ancient biological function governing the perception of chemical information to mediate essential behavioral responses related to the detection of food, pathogens, predators, conspecifics, and optimal conditions for reproduction (Benton 2015). The essential role of chemoreception at multiple levels means the molecular components of this system are key candidates for important adaptive changes associated with life in new environments. Chemical stimuli are recognised by transmembrane proteins which transform volatile and soluble substances from the environment into electrical outputs through nerve impulses. These proteins are located on the dendritic membranes of sensory neurons housed in the sensilla of receptor organs, such as antennae and other sensory appendages along the body (Joseph and Carlson 2015). In arthropods, gene families involved in chemoreception have been studied in several lineages such as the insects (Sánchez-Gracia et al. 2009), decapods (Kozma et al. 2020), and spiders (Vizueta et al. 2017). This research has pinpointed six main protein families involved in chemoreception in insects: odorant receptors (ORs), gustatory receptors (GRs), ionotropic receptors (IRs), sensory neuron membrane proteins (SNMPs/CD36), chemosensory proteins (CSPs), and odorant-binding proteins (OBPs). GRs are the oldest animal chemoreceptor family, dating back to the origin of animals (Stocker 1994; Eyun et al. 2017). In protostomes, chemical detection is also mediated by IRs, which originated as a divergent clade of ionotropic glutamate receptors (IGluRs) and N-methyl-D-aspartate receptors (NMDARs) (Benton et al. 2009). IRs have a variety of functions, such as odorant perception, thermosensation, and hygrosensation, and are principally and widely expressed in peripheral sensory systems (van Giesen and Garrity 2017). IGluRs and NMDARs are glutamate-binding excitatory neurotransmitters that play a role in brain synaptic communication (Mayer and Armstrong 2004) and regulate developmental and reproductive processes in insects (Chiang et al. 2002) respectively. Furthermore, the sensory perception of smell in insects is mediated by ORs and SNMPs. ORs are a highly diverse chemoreceptor family that participate in the detection of airborne chemical compounds, particularly in appendages on the head (Clyne et al. 1999; Robertson et al. 2003; Thoma et al. 2019), and SNMPs play key roles in chemoreception such as a highly sensitive pheromone detection (Nichols and Vogt 2008; Gomez-Diaz et al. 2016). These transmembrane proteins are assisted by low-molecular-weight soluble proteins that act in parallel as carriers of hydrophobic compounds such as some odors and tastants, thus enhancing chemical perception. These carriers are encoded by the CSP and the insect-specific OBP genes (Pelosi et al. 2014; Pelosi et al. 2018).

An extensive positive link between the extent of the chemosensory gene repertoire and the complexity of its chemical ecology has been found in insects and other arthropods occupying a large variety of ecological niches (Ngoc et al. 2016; Robertson et al. 2018; Almudi et al. 2020). In Coleoptera, remarkable expansions of odorant receptors (Mitchell et al. 2020) and other chemosensory gene families (Andersson et al. 2019) have been found in polyphagous/generalist species, whereas stenophagous/specialist species show more reduced gene repertoires. Nevertheless, the comparisons performed to date are mostly based on distantly related species undergoing completely different evolutionary histories, therefore a narrower phylogenetic context is needed to obtain more robust evidence of this correlation. Current research represents a small proportion of beetle diversity, and functional characterization of chemosensory proteins is noticeably lacking (Mitchell and Andersson 2021). Chemosensory gene families have been inferred to experience a fast dynamic evolution, where gene gain and loss are the major source of variation, providing opportunities for adaptation to new habitats (Sánchez-Gracia et al. 2009).

Caves and other subterranean environments are characterized by a key selective pressure: the permanent absence of light. Underground habitats also have a high climatic stability compared to surface habitats, with constant temperature and high humidity (Lauritzen 2018). Furthermore, nutrient inputs tend to be heterogeneously distributed and are generally scarce (Kováč 2018). All these conditions inherently influence abundance, volatility and distribution of chemical compounds and consequently may have impacted the sensory systems of its inhabitants at the phenotypic level. For example, the broadly convergent loss of eyes in cave-dwellers have been proposed to be compensated by enhanced mechanosensory and chemosensory capabilities (Poulson 1963). Sensory modifications include elongated antennae containing specific sensory organs in beetles (Accordi and Sbordoni 1977), modifications in the taste buds and lateral line in fishes (Varatharasan et al. 2009; Yoshizawa et al. 2014), and enhanced development of olfactory brain regions in crustaceans (Stegner et al. 2015). Nevertheless, very little is known about the molecular evolution of the chemosensory system in cave-dwelling fauna, particularly from a macroevolutionary perspective, with only a few studies so far focusing on single species at a time (Yang et al. 2016; Balart-García et al. 2021). Previous studies on the evolution of the chemosensory capabilities of Coleoptera determined that the odorant and gustatory capabilities of a cave beetle were relatively reduced and seemed to follow a similar pattern than other beetle specialists (Balart-García et al. 2021). Assuming that larger OR and GR repertoires are correlated with broader capabilities to perceive different odorant and gustatory stimuli, the results suggested that cave beetles only differentiated a relatively poor diversity of chemical compounds. This result fitted well with the fact that the subterranean environment is relatively less complex in terms of chemical information compared to surface habitats due to its particular conditions such as poor nutrient diversity, environmental homogeneity and less biodiversity. Perhaps cave beetles have an extremely sensitive smelling and tasting system facilitating the perception of particular chemical stimuli in the darkness, but according to this study the diversity of odorants and tastants they can perceive and interpret is reduced compared to surface-dwelling species. Extreme sensitivity to detect food items in the darkness was previously observed in the behavior of a highly specialized cave beetle of the family Carabidae, which predate on cave-cricket eggs and display targeted movements to distant and hidden oviposition sites (Kane and Poulson 1976). Leiodidae cave beetles have also shown olfaction responses when baiting caves with different food items (Peck, 1984). The scavenger and saprophagous species, such as the species of the tribe Leptodirini, tend to forage by ‘constant walking’ covering large areas to find the patchily distributed food items and often aggregate when there is a substantial food input from the surface (PBG, pers. observation).

Cave beetles constitute an excellent system to understand the evolutionary processes leading to adaptation to subterranean habitats. Beetles comprise numerous instances of phylogenetically-distinct subterranean lineages with varying degrees of morphological change and ecological specialization related to their adaptation to underground habitats. The tribe Leptodirini (Leiodidae, Cholevinae) represents the largest animal radiation in subterranean habitats and it is principally distributed in the North of the Mediterranean basin. This beetle tribe represents ancient colonizations of underground habitats (ca. 30 Mya) including multiple lineages with a varying degree of specialization to life in caves. Virtually all the species of the tribe Leptodirini are cave-dwelling or live in forest litter, being morphologically similar to cave-dwelling species (Ribera et al. 2010; Fresneda et al. 2011). Furthermore, diving beetles (Dytiscidae, Hydroporinae) also show multiple independent instances of subterranean colonization by surface species of isolated calcrete (carbonate) aquifers in arid regions of Western Australia, especially in the tribes Bidessini and Hydroporini (Cooper et al. 2002; Leys et al. 2003). These aquatic beetle tribes underwent underground colonization more recently (ca. 7–3 Mya) and show clear phenotypic differences between surface-dwelling and subterranean species (Leijs et al. 2012; Langille et al. 2021; Zhao et al. 2023). The chemosensory gene repertoire has been characterized so far in a single cave species, the Leptodirini beetle *Speonomus longicornis*. This species presents a diminished diversity of odorant and gustatory gene repertoires compared to polyphagous beetles inhabiting surface habitats, and contains a gene duplication of the ionotropic coreceptor IR25a, a highly conserved single-copy gene in protostomes (Balart-García et al. 2021) involved in thermal and humidity sensing among other olfactory roles (Ni 2020). Nevertheless, the macroevolutionary processes governing chemosensory gene family evolution across cave-dwelling taxa have never been studied.

Here, we investigate the molecular evolution of the chemosensory system in beetles that have undergone independent instances of subterranean colonization using a genome-wide phylogenomic approach. Specifically, we annotated the main chemosensory gene families and characterized gene repertoire evolutionary dynamics in beetle species with a broad range of habitat preferences (i.e., aquatic, terrestrial, surface-dwelling and subterranean ecologies) to understand the role of gene gain, duplication and loss in the evolution of gene families involved in chemosensation. At least six underground transitions are represented by lineages of the tribes Leptodirini (terrestrial) and Bidessini and Hydroporini (aquatic), which allowed us to explore how the chemosensory gene repertoire was reshaped in the context of parallel subterranean evolution. Our study sheds light into the molecular evolution of the chemosensory systems of cave-dwelling fauna from a macroevolutionary perspective.

## 2. MATERIALS AND METHODS

### 2.1 Data collection, chemosensory gene families characterization and phylogenetic inferences

We used the same dataset as the one used in Balart-García et al. (2023) to explore gene repertoire evolution, consisting of highly complete transcriptomes and genomes for a total of 39 Coleoptera, one Strepsiptera and one Neuroptera species. The studied Coleoptera inhabit aquatic and terrestrial habitats, both including surface-dwelling and subterranean ecologies. The aquatic lineage (Dytiscidae, Hydroporinae) include two species from the tribe Hydroporini and six species from the tribe Bidessini. The terrestrial lineage (Leiodidae, Cholevinae), include a total of 13 species from the tribe Leptodirini and one species from the tribe Catopini (i.e., *Catops fuliginosus*) representing a closely related outgroup. We annotated genes encoding chemosensory related proteins (i.e., OR, GR, IR, IGLUR, NMDAR, SNMP, OBP and CSP) for the 41 proteomes by using BITACORA v.1.3 (Vizueta et al. 2020) in “protein mode”. This sequence similarity based software combines BLAST and HMMER searches using custom databases that contain sequences of the gene families of interest. We generated curated databases for each chemosensory gene family which include sequences of *Drosophila melanogaster* (flybase.org), that had been experimentally confirmed (i.e., GRs, IGluRs, NMDARs and SNMP/CD36s), and sequences of several Coleoptera species that were characterized in previous studies using comparative genomics methods (ORs, GRs, IRs, SNMPs, CSPs, OBPs) (Dippel et al. 2014, 2016; Schoville et al. 2018; Andersson et al. 2019; Zhao et al. 2020). Due to the high heterogeneity of some chemosensory gene families and the inclusion of partial genes in our proteome sets, the dubious hits obtained with each database (e.g., result significant both for OR and GR hits or IGluR and IR hits) were identified and manually validated using additional protein searches with InterPro (Blum et al. 2021). Proteomes of species whose chemosensory gene families were characterized in Andersson et al. (2019) and Mitchell et al. (2020) (i.e., *Tribolium castaneum*, *Agrilus planipennis*, *Onthophagus taurus*, *Nicrophorus vespilloides*, *Anoplophora glabripennis* and *Dendroctonus ponderosae*) were re-annotated with our approach in order to follow the same methodology for all the data sets, therefore reducing the potential biases arising from the different annotation methods used in different studies.

We conducted maximum-likelihood phylogenetic inferences for each chemosensory gene family in order (i) to validate the annotation of the candidate chemosensory gene sets and discard dubious sequences, (ii) to identify highly conserved chemosensory subclades across insects, and (iii) to study their phylogenetic relationships across Coleoptera. To do this, we included some sequences of the curated databases in the phylogenetic inferences as references for the main chemosensory gene family clades. These references consisted of GR, IGluR, NMDAR, SNMP/CD36 sequences of *D. melanogaster*, OR, GR, IR, CSP, SNMP/CD36 and OBP sequences of *T. castaneum,* GR sequences of *A. planipennis* and *D. ponderosae* and a variety of Coleopteran SNMP sequences obtained from Zhao et al. (2020). We aligned the amino acid sequences for each chemosensory gene family using MAFFT v.7.407 (Katoh and Standley 2013) with a maximum of 1,000 iterations. We trimmed the alignments with trimAl v.1.4.1 (Capella-Gutiérrez et al. 2009) using the option –gt 0.4 and manually discarded sequences with less than 10% of the total amino acid positions in the alignment. We used FastTree v.2.1.11 under the LG model (Price et al. 2010) to generate a gene tree that was then implemented as a guide tree for the final phylogenetic inference with IQ-TREE v.2.1.3 (Minh et al. 2020) using the mixture model LG+C20+F+G with the site-specific posterior mean frequency model (PMSF; Wang et al. 2018) and the ultra-fast bootstrap option (Hoang et al. 2018). We used iTOL v6 to visualize and render the phylogenetic trees (Letunic and Bork 2021).

### 2.2 Estimation of the evolutionary dynamics of the chemosensory gene repertoire

We used BadiRate (Librado et al. 2012) to study the evolutionary dynamics of the chemosensory gene families under the birth, death and innovation model (BDI). This software uses a maximum-likelihood based approach to estimate gene gains and losses across all branches in a given phylogenetic context. We performed this estimation for each chemosensory gene family by using the candidate gene counts (Supplementary Table 1) and the time-calibrated tree that was generated in Balart-García et al. (2023). Furthermore, to investigate the most relevant evolutionary events that could lead to parallel significant changes in the chemosensory gene families, we examined significant changes in the net rate (i.e., the difference between gene gain and loss rates, henceforth ‘net rate’) under different branch models. We compared a “global rates” (G) estimation model— which assumes that all of the branches have the same net rate—to several “branch-specific” rates models, that assume that a certain set of selected branches have different net rates compared to the background. These branch-specific models consisted of (i) the branches of the most recent common ancestors of the tribes (MRCA model), (ii) the branches leading to the underground habitat shift (HS model) and (iii) all the branches of subterranean lineages together (SB model). These analysis were performed including all the taxa, but in order to explore lineage-specific shifts in the chemosensory gene families, we performed the analysis at a narrower phylogenetic context either with only the aquatic tribes (i.e., Bidessini and Hydroporini) or the terrestrial tribe (i.e., Leptodirini) separately. For the terrestrial tribe analysis we performed an extra branch-specific model which assumes a rate shift in the branches of the highly modified lineages (HML) compared to the background and thus allowing us to explore whether remarkable changes in the chemosensory repertoire occurred within a subterranean ecosystem. These highly modified lineages correspond to two clades of the tribe Leptodirini that parallelly developed a life cycle contraction and extremely modified body shape and elongation of sensory appendages (i.e., *Speonomus longicornis* – *Troglocharinus ferreri*; *Astagobius angustatus* – *Leptodirus hochenwartii*) (Deleurance-Glaçon 1963; Delay 1978; Cieslak et al. 2014). Finally we calculated the Akaike information criterion (AIC) and compared the global rates model to each branch-specific model independently at every phylogenetic level (i.e., all taxa, aquatic lineage, terrestrial lineage) in order to find the best fitting model.

## 3. THEORY

Radical ecological shifts, such as subterranean colonization, could have a strong impact on the extent and composition of the chemosensory capabilities, and could lead to parallel genetic changes across distantly related lineages. In a previous study, the cave-dwelling beetle *Speonomus lognicornis* showed a relatively reduced repertoire of some chemosensory gene families (i.e., odorant receptors and gustatory receptors) compared to distantly related Coleoptera species with a wide range of ecological preferences (Balart-García et al. 2021). Some highly conserved genes were found to be duplicated or lost in this species (e.g., the ionotropic receptor 25 and gustatory receptors related to sugar detection respectively). Nevertheless, more comparisons in a narrower phylogenetic context were needed to explore the extent of these genetic changes in strictly subterranean beetles and the degree of parallel evolution of their chemosensory system. In this piece of work we expanded the phylogenetic framework including multiple lineages of terrestrial (i.e., tribe Leptodirini) and aquatic (i.e., tribes Bidessini and Hydroporini) Coleoptera that independently transitioned to caves and compared the chemosensory gene repertoires of surface-dwelling and subterranean lineages. This exploratory approach aims to shed light on candidate chemosensory gene duplications and losses in cave-dwelling beetles and to further our understanding of the evolution of insect chemosensory gene families from a macroevolutionary perspective.

## 4. RESULTS AND DISCUSSION

### 4.1 Highly heterogeneous chemosensory gene repertoires in Coleoptera are potentially driven by diverse ecological preferences

Our data set includes 22 transcriptomes for surface-dwelling and subterranean species of the tribes Bidessini, Hydroporini, and Leptodirini including at least six independent beetle lineages that colonized subterranean habitats. Moreover, we annotated candidate chemosensory genes for a total of 39 Coleoptera, one Strepsiptera and one Neuroptera species based on highly complete proteomes (Balart-García et al. 2023). Our results indicate a high variability in the extension and composition of the chemosensory gene repertoires across Coleoptera (Figure 1, Supplementary Table 1). ORs and OBPs are the most variable chemosensory gene families in terms of copy number variation (i.e., standard deviations of ORs and OBPs are 32,4 and 25,3 respectively, followed by GRs (24), IRs (12.3), CSPs (4.3), IGluRs (4), SNMPs (2.6) and NMDARs (1.9)). This result suggests that these two insect-specific chemosensory gene families, related to odorant perception, have undergone a more dynamic evolution compared to more ancient gene families that resulted in relatively less variable repertoires. Noticeably, species within Adephaga show more reduced chemosensory gene repertoires than the Polyphaga (i.e., with 26.6 and 87.5 of standard deviation respectively).

**Figure 1.**
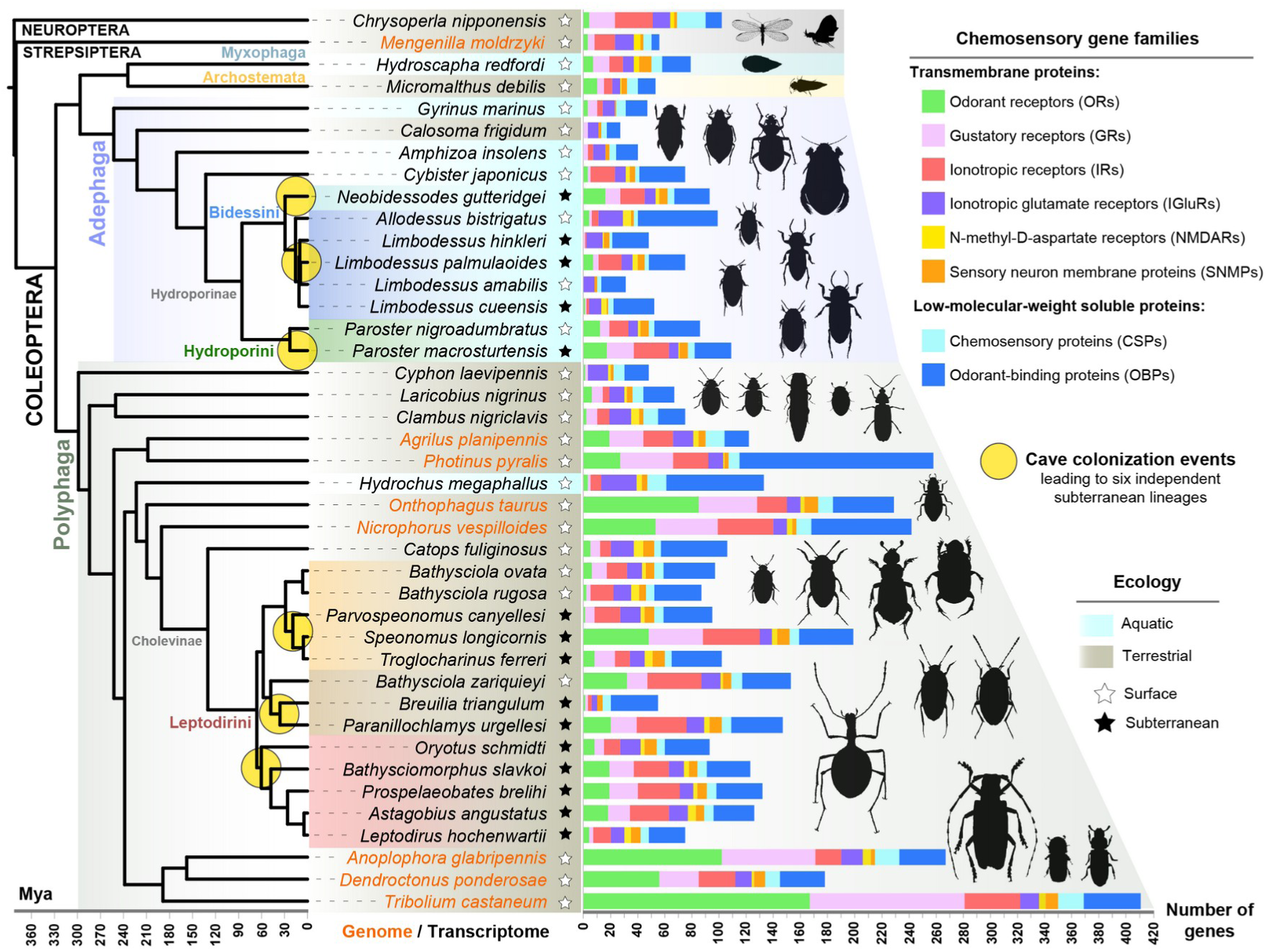
Phylogenetic context and size of the chemosensory gene repertoires in Coleoptera. Time-calibrated phylogeny of the studied Coleoptera and outgroups obtained from Balart-García et al. (2023), including six cave colonization events leading to independent subterranean lineages: two in the tribe Bidessini (light blue and dark blue), one in Hydroporini (green) and three in Leptodirini (orange, brown and red). Orange tips indicate genomes and black tips indicate transcriptomes. The central column indicates the ecology of the studied species (i.e., aquatic – terrestrial, surface – subterranean). Bar plots indicate the total number of chemosensory genes annotated for each species.

Large expansions in olfactory and gustatory related gene families may be linked to the perception of complex and heterogeneous environments in terms of chemical stimuli (e.g., Andersson et al. 2019, Ngoc et al. 2016), which appear to be frequent in terrestrial ecosystems (i.e., more prevalent in Polyphaga). Polyphaga represents the coleopteran suborder with the most diverse feeding habits (Grimaldi & Engel 2005), including predators, scavengers, fungal grazers, and feeders of algae, bacteria and macrophytes. Conversely, the Adephaga species we studied are mostly aquatic predators and they might not principally rely on airborne chemical clues to navigate in their environment. Furthermore, the early branching lineages of Polyphaga, represented by *Cyphon laevipennis* (Scirtidae), retain some ancestral habitat preferences (i.e., aquatic or semi aquatic larvae; Lawrence 2016) and show a reduced chemosensory gene repertoire (i.e., 48 genes), whereas other polyphagan species show remarkable expansions in specific gene families. The latter includes the firefly *Photinus pyralis* with 165 OBPs, and the red flour beetle and Asian longhorn beetle (*Tribolium castaneum* and *Anoplophora glabripennis*), which show extremely large gene repertoires that may reflect their extended chemosensory capabilities (e.g., 433 and 289 annotated genes respectively). However, proteome data from these two species were based on whole genome sequence data, and it is likely that transcriptome data, despite obtaining high completeness scores based on BUSCO databases, could have provided an underestimate of the actual number of chemosensory genes in each family. Nevertheless, in comparisons of transcriptome data, the only terrestrial adephagan species that we studied (i.e., *Calosoma frigidum*; 27 genes) shows a very reduced chemosensory gene repertoire and, in contrast, the aquatic polyphagan species (i.e., *Hydrochus megaphallus*; 133 genes) shows a relatively large chemosensory gene repertoire. In essence, these results suggest that the extent of the chemosensory gene repertoires in Coleoptera might be strongly related to the habitat preferences and feeding habits, where aquatic predaceous beetles show relatively moderate chemosensory capabilities compared to terrestrial polyphagous beetles. Further studies, including complete genomes of Adephaga species and more representatives of terrestrial and aquatic ecologies of these two suborders, would help to obtain more robust evidence of these contrasting trends.

The sizes of the chemosensory repertoires in beetle tribes that have colonized underground habitats (i.e., Bidessini, Hydroporini and Leptodirini) do not show a clear pattern when comparing surface-dwelling and subterranean species. The subterranean species of Hydroporini (i.e., *Paroster macrosturtensis*) shows 26 more chemosensory genes than its surface-dwelling relative (i.e., *Paroster nigroadumbratus*) and the subterranean Bidessini species of the genus *Limbodessus* (i.e., *L. hinkleri*, *L. cueensis* and *L. palmulaoides, with 52, 48 and 75 annotated genes respectively)* show larger chemosensory gene repertoires than their corresponding surface-dwelling relative within the same genera (i.e., only 31 annotated genes in *L. amabilis*) (Supplementary table 1). These chemosensory gene repertoire expansions in subterranean species of the genus *Paroster* and *Limbodessus* suggest that the subterranean environment might lead to chemosensory gene family expansions that provide broader capabilities in the perception of chemical stimuli. This change could be related to the increased scavenging habits of subterranean species compared to the predatory preferences of their surface-dwelling relatives, or a dietary shift from predation to omnivory that broadly converges in many subterranean taxa (Gibert and Deharveng 2002, Hüppop 2012). Nonetheless, a surface species, *Allodessus bistrigatus, which is closely related to Limbodessus taxa,* shows an opposite pattern with an expanded suite of chemosensory genes, including a large expansion in the OBP family (i.e., 56 OBPs and a total of 99 annotated genes). Further comparisons, including surface-dwelling and subterranean relatives for each genus, need to be made to confirm the previously-mentioned pattern of enhanced chemosensory capabilities in the subterranean aquatic beetles.

In the tribe Leptodirini, we observe a contrasting pattern between two subterranean species: *Speonomus longicornis* has the largest chemosensory gene repertoire, whereas *Breuilia trianglum* exhibits a very reduced one (199 and 55 annotated genes respectively; Figure 1 and Supplementary Table 1). The large repertoire of chemosensory genes observed in *S. longicornis*, particularly for odorant receptors, could be explained by the inclusion of separate antennae samples and thus a relatively more exhaustive protocol for characterizing chemosensory genes (i.e., sequencing mRNA from structures with chemosensory specificity). Nevertheless, when comparing closely related cave-dwelling species, whose genetic repertoires were characterized from exactly the same sample types, transcriptome sequencing methods and obtaining similar completeness scores, such as *Leptodirus hochenwartii* and *Astagobius angustatus*, we observed a large variation in their chemosensory repertoires (i.e., 75 and 126 annotated genes respectively). This result suggests that the chemosensory gene families have undergone extensive changes at the lineage-specific level in strictly subterranean species. Therefore, the chemosensory gene repertoire varies dramatically between species, and there seems to be no pattern with general trends when comparing surface to cave-dwelling species. Both surface-dwelling and subterranean species of the tribe Leptodirini are mainly detritivorous or saprophagous, feeding on carrion, fungi and biofilms. However, some cave-dwelling species show modifications of the mouthparts associated with a highly specific dietary niche. For instance, some subterranean species are adapted to a semiaquatic lifestyle, such as found in hygropetric habitats on cave walls, and filter organic particles with their modified mouthpieces (Moldovan 2004). Lineage-specific dietary specializations in Leptoridini may have contributed to the highly diverse chemosensory gene repertoires that we observed in surface and cave-dwelling species.

### 4.2 Rapid and complex evolution of the chemosensory system in subterranean beetle lineages

We used maximum likelihood-based phylogenetic comparative methods to explore parallel evolution in the chemosensory gene repertoires of subterranean lineages. We investigated significant gene gain and loss shifts for each chemosensory gene family (i.e., the difference between gene gain and loss rates, net rate hereafter) prior, during, and after the subterranean transition. None of the branch-specific rate models were significantly better-fitting compared to the global rate model for any chemosensory gene family, indicating no parallel changes in the net rate of evolution of chemosensory gene families occurred prior, during, or after the subterranean transition in the selected tribes (Supplementary Table 2). Moreover, we could not obtain the net rate estimation for the odorant receptors (OR) and gustatory receptors (GR) under any branch-specific rates model due to the high heterogeneity of the gene counts across the selected taxa and the complexity of the branch-specific models. At the lineage-specific level, we obtained the net rate estimations under all the models and it was possible to estimate the rate of evolution of ORs. In the Leptodirini-specific analysis, ORs resulted in a significantly better fit under the habitat shift (HS) model, but only after excluding the outlier *S. longicornis.* Our results thus indicate a contraction of the OR repertoire when lineages transitioned underground in Leptodirini. Nonetheless, given the outlier and the limited representation of surface-dwelling species, the evolution of odorant receptors in cave-dwelling fauna need to be further investigated to obtain more robust evidence.

We also investigated the estimated gene gains and losses per branch for each chemosensory gene family using the results of the global rates model. This allowed us to explore in detail where the most relevant pulses of chemosensory gene repertoire change occurred in the phylogenetic context of this study (Figure 2). Our results indicate that gene family expansion was more relevant than contraction in most of the species. Nevertheless, we found some species that primarily showed gene family contractions (i.e., *Limbodessus amabilis*, *Troglocharinus ferreri*, *Breuilia triangulum* and *Leptodirus hochenwartii).* Despite not being significant under the previously mentioned MRCA branch-specific model, when dissecting the ancestral reconstruction of gene gains and losses we found some gene families experiencing parallel changes in the MRCA of each tribe. For instance, OBPs and CSPs were expanded in parallel, and IGluRs were parallelly contracted in the MRCA of the three tribes. On the other hand, the IR family was contracted in the MRCA of Leptodirini and Bidessini, and expanded in Hydroporini. The SNMP family was only expanded in the MRCA of Leptodirini and Hydroporini. These parallel and exclusive changes could represent examples of genomic exaptation that potentially led to chemosensory adaptation in underground habitats. Moreover, we did not observe substantial changes in the branches where species transitioned underground, suggesting that the chemosensory families experienced lineage-specific changes within surface and subterranean lineages. More remarkably, and despite not being significant in the branch-specific model, the ancestral estimations of gene gain and loss in the highly modified lineages (HML) of Leptodirini (i.e., with a single larval-instar life cycle and presenting extremely derived subterranean phenotypes) indicate a parallel expansion of GRs, IRs, and, to a lesser extent, an expansion of NMDARs. These results suggest highly specialized subterranean lineages that colonized caves >30 Mya, despite not having undergone a substantial reshaping in the chemosensory gene repertoire, could have experienced key parallel changes in some chemosensory gene families. When further exploring the expansions and contractions in these highly modified species we observed contrasting patterns, in which *A. angustatus* and *S. longicornis* show large expansions of GRs and IRs compared to the gene family dynamics of *L. hochenwartii* and *T. ferreri,* that essentially consist of contractions. Likewise, the lineage leading to the surface-dwelling species *Bathysciola ovata* + *Bathysciola rugosa* also experienced a relevant expansion in several chemosensory gene families. Altogether, these results indicate that chemosensory gene families experienced rapid evolution including expansion and contractions in surface and subterranean species. Contrasting patterns in closely related species suggest that no remarkable quantitative changes occurred due to subterranean colonization, and thus that changes occurred independently within subterranean ecosystems.

**Figure 2.**
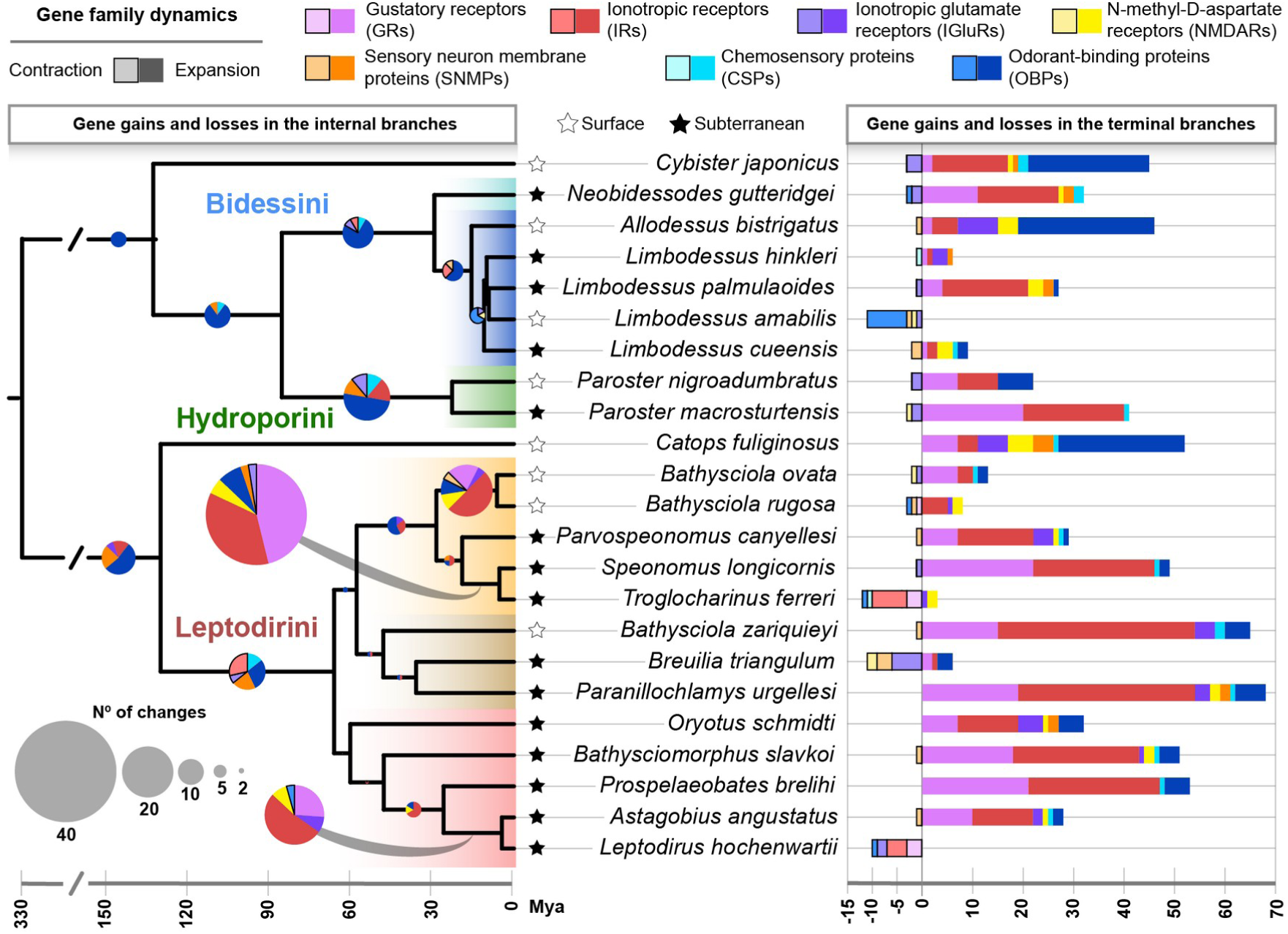
Ancestral contractions and expansions of the chemosensory gene families using a likelihood-based estimation with BadiRate. Time-calibrated tree indicating the total number of gene gains and losses of each chemosensory gene family in the internal branches (pie charts) and in the terminal branches (bar plots). Pie charts and bar plots are scaled to the total number of gene gains and losses of each chemosensory gene family. The different colored ranges at the species tree represent the six independent instances of underground evolution, with black stars indicating subterranean species and white stars surface-dwelling species of each clade.

The odorant receptor (OR) gene family was significantly contracted in the subterranean lineages of Leptodirini, with the exception of *S. longicornis* that showed the most extended chemosensory gene repertoire within Leptodirini (Figure 3). Furthermore, our analyses revealed a low number of genes in this gene family in the MRCA of each clade and a progressive expansion towards the tips, suggesting that a fast dynamic of expansions occurred at a lineage-specific level and was the driving process guiding the evolution of ORs.

**Figure 3.**
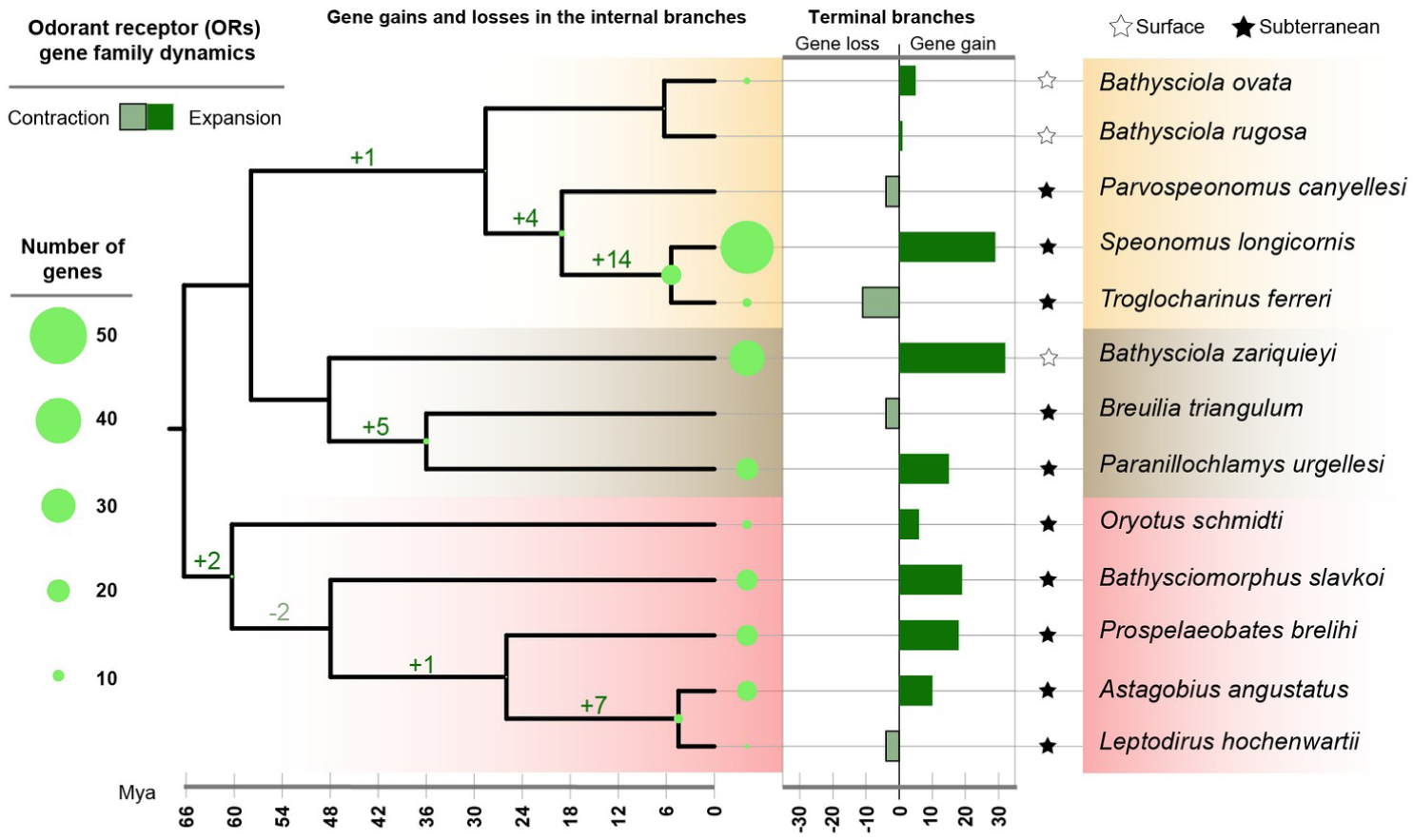
Odorant receptors (ORs) evolution in the tribe Leptodirini based on a likelihood estimation with BadiRate. Time-calibrated phylogeny indicating the estimated size of the OR repertoires (green dots) and the number of gene gains and losses in the internal branches. Barplots indicate the proportion of gene gain and loss in the terminal branches. White stars indicate surface-dwelling species, black stars indicate subterranean species and colored circles indicate lineages representing independent underground colonization events.

### 4.3 Parallel gene gains and losses in key chemosensory gene families may have facilitated the adaptation to life in caves

Most of the conserved chemosensory clades appear highly supported in our phylogenetic inferences (Figure 4; see also Supplementary Figures 1–7). However, there are some differences in the relationships between some clades compared to those obtained in previous studies that included a more reduced set of Coleoptera species. For instance, the OR2A clade was not a monophyletic group in our analysis and members of this clade, previously reported in *T. castaneum,* were placed into two supported subclades (Figure 4). Moreover, we detected nine highly-supported CSP clades for Coleoptera (i.e., A–I) and one clade composed only by Neuroptera and Strepsiptera sequences (i.e., O). Some of these CSP clades correspond to lineage-specific expansions, for instance the clade I is specific to *Tribolium castaneum*, H is specific to Tenebrionoidea + Chrysomeloidea + Curculionoidea and E is specific to Hydrophiloidea + Scarabaeoidea + Staphylinoidea (Supplementary Figure 1). Regarding the OBP clades, we found two supported clades for the antennal binding proteins II (ABPII) and for Classic OBPs and four clades of Minus-C OBPs. Interestingly, most of the OBPs of *Photinus pyralis*, which would represent one of the largest OBP repertoires reported in insects, correspond to divergent OBPs with multiple duplications (Supplementary Figure 2). Nonetheless, we were not able to characterize any OR, GR and some of the conserved IR clades in some species (Supplementary Table 1), indicating that a more exhaustive approach needs to be used to clarify the evolution of these transmembrane chemoreceptor gene families in Coleoptera. Gene expression data from different chemosensory appendages showing functional specificity, such as antennae and mouthparts, and comparisons with highly complete genomes of the same species would also help to characterize in detail these chemoreceptor families.

**Figure 4:**
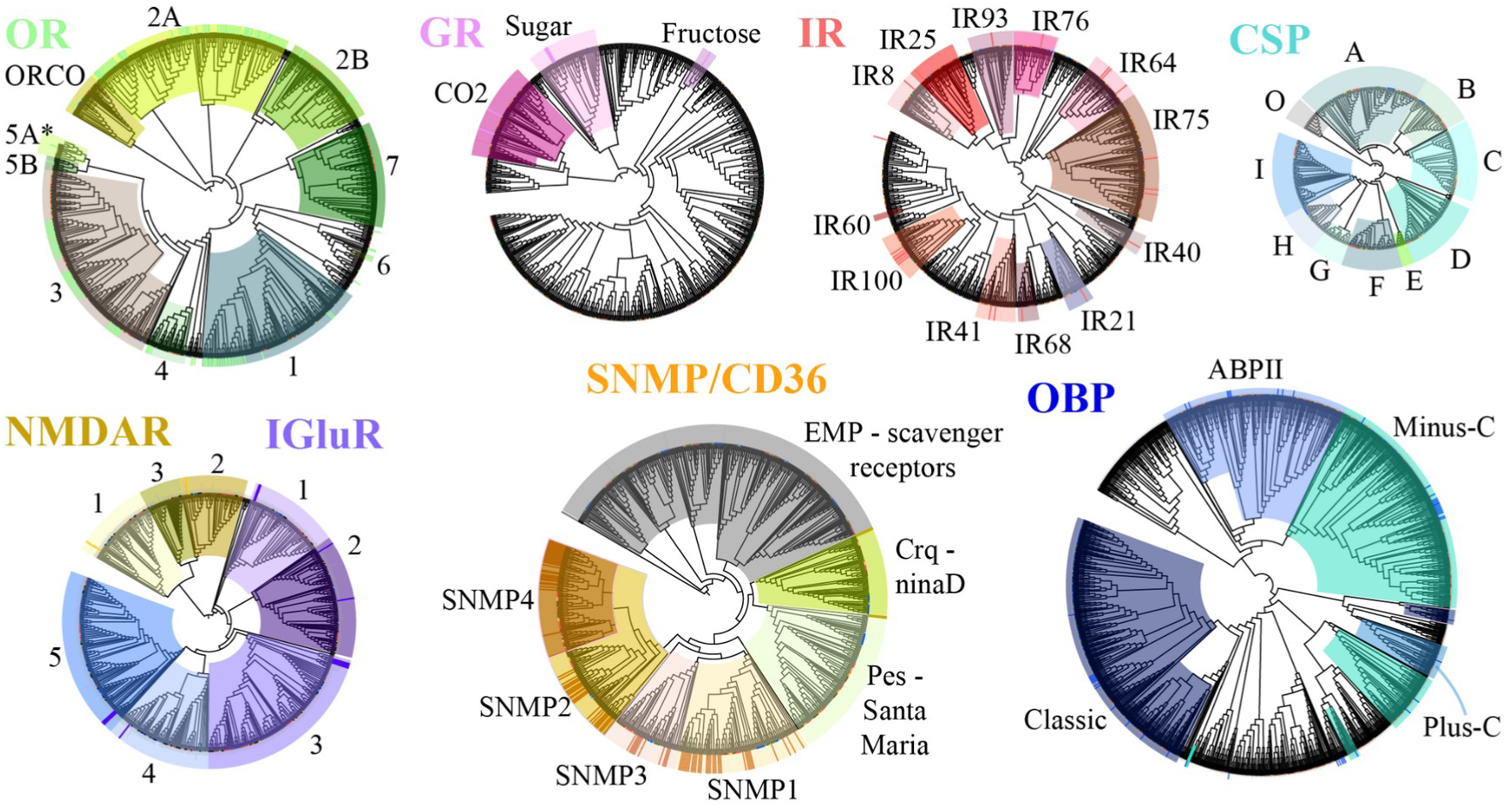
Chemosensory gene families and supported clades in Coleoptera. Maximum-likelihood phylogenetic trees for each chemosensory gene family as follows: odorant receptors (OR), gustatory receptors (GR), ionotropic receptors (IR), N-methyl-D-aspartate receptors (NMDAR), ionotropic glutamate receptors (IGluR), sensory neuron membrane proteins (SNMP/CD36), chemosensory proteins (CSP) and odorant-binding proteins (OBP). Colored ranges indicate significantly supported clades (i.e., bootstrap >75%) identified with reference sequences from a variety of studies (i.e., indicated with highlighted outer strips, see Methods). The lineage-specific OR5A (*) clades of *Tribolium castaneum* are collapsed (i.e., 227 ORs). See Supplementary Figures 1-7 for more detailed information.

The conserved gustatory receptors related to the perception of sugar and fructose were not detected in any Leptodirini species nor in *Catops fuliginous* (both in the same subfamily, Cholevinae) (Figure 5A). However, *Nicrophorus vespilloides*, the closest relative to Cholevinae in our study (i.e., same superfamily, Staphylinoidea) showed three gene copies of candidate sugar receptors. Our results thus indicate that the gustatory perception of sugar and fructose has potentially been lost in Cholevinae. On the other hand, we detected sugar receptors in two subterranean species of the tribes Hydroporini and Bidessini (*Paroster macrosturtensis* and *Limbodessus palmulaoides*), indicating that the aquatic clade retains sugar perception capabilities. Furthermore, the apparently conserved fructose receptor clade previously characterized in some phytophagous Polyphaga species is also present in *Laricobius nigrinus, a* predatory specialist of the hemlock woolly adelgid, indicating that this fructose receptor clade is not exclusive to species merely feeding on plants, but is also conserved in species whose diet may include substantial inputs of fructose (Cohen and Cheah 2015).

**Figure 5.**
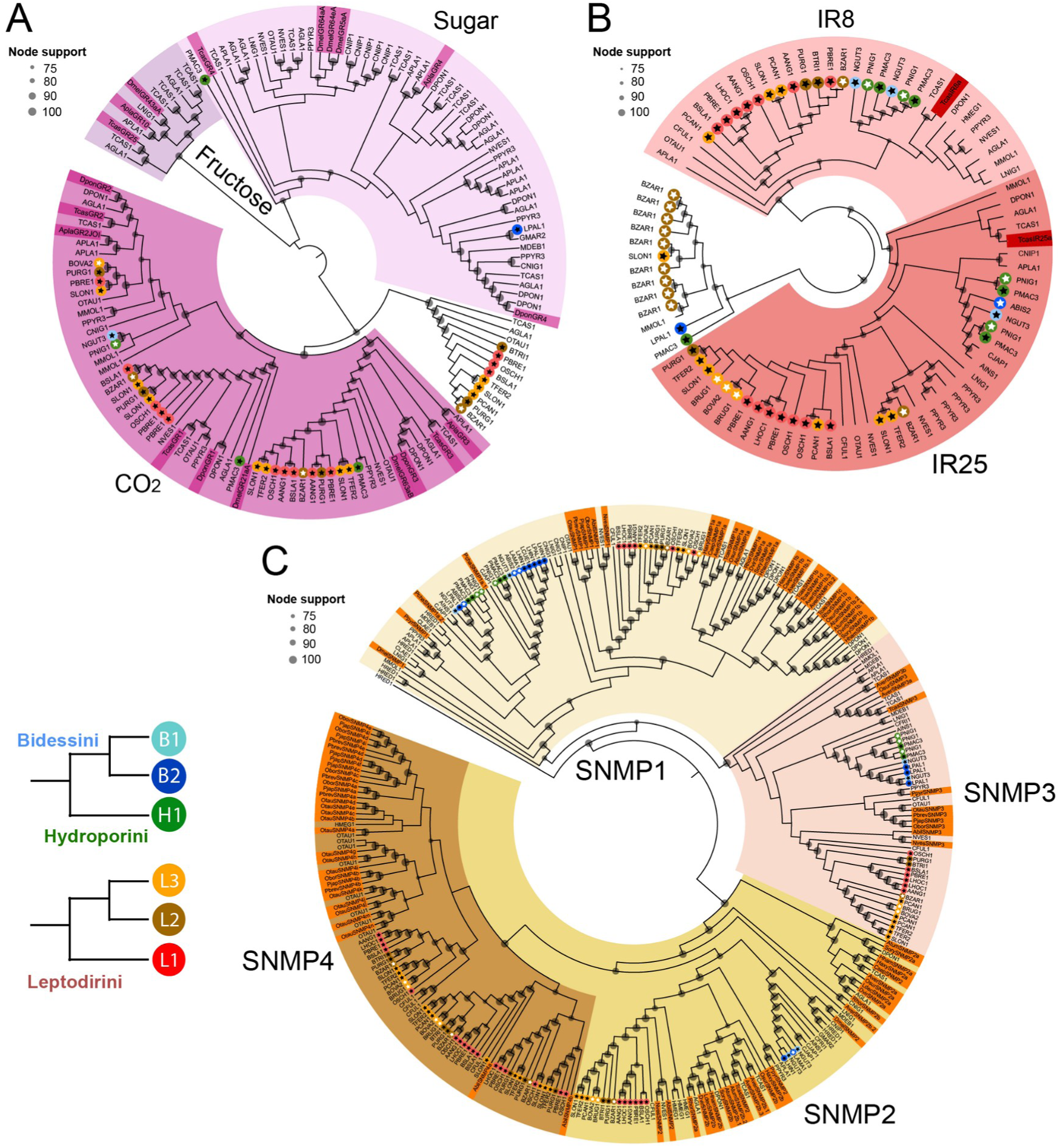
Conserved gustatory (GR) and ionotropic (IR) receptor clades and sensory neuron membrane protein (SNMP) clades in Coleoptera. Pruned maximum-likelihood gene trees of GR (A), IR (B) and SNMP (C). White stars represent surface-dwelling species and black stars subterranean species. The highlighted tips correspond to previously annotated chemosensory genes used as a reference to identify the main clades (see Methods). For the complete gene trees see Supplementary Figures 4, 5 and 6.

We further explored the conserved clade of CO_2_ gustatory receptors, which typically consist of two or three gene copies in Coleoptera (Andersson et al. 2019). Two subterranean species of the tribe Leptodirini (i.e., *Speonomus longicornis* and *Prospelaeobates brelihi*) showed more than three copies of these highly conserved CO_2_ receptors (Figure 5A, Supplementary Figure 4), suggesting parallel gene duplications of these CO_2_ receptors in independent subterranean lineages. Remaining Leptodirini species showed a variable number of CO_2_ receptors (from zero to three). On the other hand, very few CO _2_ receptors were identified in Hydroporini (i.e., one copy in the surface-dwelling species and two copies in the subterranean species), and just one copy in a surface-dwelling species of Bidessini. Moreover, we detected a highly supported sister clade of these CO_2_ receptors consisting of sequences of primarily Leptodirini species, but also other Polyphaga species. In summary, these results indicate that CO_2_ receptors are less conserved than originally suggested in previous studies and that possibly the species with absent receptors may perceive CO_2_ via other kinds of receptors.

Ionotropic receptors (IR), highly conserved in virtually all protostomes, also showed some novelties in the studied beetle tribes. A surface-dwelling species and several subterranean species of the tribe Leptodirini (*Bathysciola rugosa*, *Speonomus longicornis*, *Troglocharinus ferreri*, *Prospelaeobates brelihi*, *Oryotus schmidti*) and Hydroporini (*Paroster macrosturtensis* and *Paroster nigroadumbratus*) showed candidate gene duplications of the IR25a gene, suggesting a parallel duplication of this gene not only in independent subterranean lineages, but also in their surface-dwelling relatives (Figure 5B, Supplementary Figure 5). Interestingly, the IR25a gene has been shown to be related to temperature and humidity sensing in *D. melanogaster* among other olfactory roles (Knecht et al. 2017) potentially suggesting this candidate gene duplication enhanced the perception of key environmental variables in surface and subterranean species of both Leptodirini and Hydroporini. Despite the Hydroporini species being aquatic and inhabiting environments with a highly constant temperature, *Paroster nigroadumbratus* inhabit temporary waters that are impacted by droughts and floods on the surface (Watts and Leijs 2008), thus a fine hygrosensory and thermosensory system might represent a clear advantage to detect seasonality. Surface-dwelling and subterranean species of the Leptodirini tribe inhabit environments with a high environmental humidity and constant temperature and they show a low desiccation resistance, thus detecting small variations in temperature and humidity may facilitate them to navigate and find the optimal conditions within their particular habitat. The gene duplication of IR25a has been described previously in *S. longicornis* (Balart-García et al. 2021), and in this study we expand the observation of several cave beetle species and their surface relatives. In addition, a candidate duplication was also detected in the IR8a (i.e., a basic co-receptor for odorant reception of multiple compounds together with other divergent IRs) for both surface-dwelling and cave species of Hydroporini and for two subterranean species of Leptodirini (*A. angustatus* and *P. brelihi*). A duplication of IR8a was only previously observed in copepods (Eyun et al. 2017), indicating that these cases in Leptodirini and Hydroporini species represent the first occurrence in insects.

We also detected several copies of the recently described beetle-specific clade SNMP4 found in some scarabaeid and staphylinid species (Zhao et al. 2020) (Figure 5C, Supplementary Figure 6). Our results revealed the presence of SNMP4 in *Hydrochus megaphallus* (Hydrophiloidea) and multiple copies were detected in species of the tribe Leptodirini and in *Catops fuliginosus,* the latter clustering together with the genes of their closest relative within Staphylinoidea (*Aleochara bilineata*). This finding indicates that the SNMP4, which essentially consists of a SNMP2 subclade, is likely to be specific to Hydrophiloidea, Scarabaeoidea and Staphylinoidea.

## 5. CONCLUSIONS

Here, we show that the evolution of the chemosensory gene repertoire is highly dynamic across terrestrial and aquatic cave beetle lineages, with no common pattern of substantial variation among the six independent subterranean transitions. In addition, our results indicate lineage-specific expansions and contractions in both surface-dwelling and subterranean lineages (Figure 2). Moreover, gene gain was found as a major force driving the evolution of gene families involved in chemoreception in some lineages, suggesting that in some cases the chemosensory system increased in complexity probably to compensate for the lack of information through the visual system. Key specific gene duplications and losses seem to have occurred prior to underground transitions, epitomized by the duplication of the genes IR25a and IR8a (involved in thermal and humidity sensing among other olfactory roles) in cave lineages and their surface sister species (Figure 3B), or the loss of sugar and fructose gustatory receptors (Figure 3A) prior to the diversification of the tribe Leptodirini that comprises several cave lineages. These specific duplications and losses may represent potential drivers of cave colonization, but further replication and experimental validation are required to confirm this. Overall, our findings pave the way to a deeper comprehension of macroevolutionary trends that underlie adaptation to life underground.

## 6. DATA AVAILABILITY STATEMENT

Raw reads have been deposited in the National Center for Biotechnology Information (NCBI; BioProject accession number PRJNA902350). All processed data and custom scripts have been deposited in github: https://github.com/MetazoaPhylogenomicsLab/Balart_Garcia_et_al_2023_chemosensory_gene_repertoire_cave_beetles

## ACKNOWLEDGEMENTS

PBG was supported by an FPI grant (grant agreement no. BES-2017-081050) financed by MCIN/AEI /10.13039/501100011033 and by European Social Fund (ESF) ‘Investing in your Future’ and the Systematics Research Fund 2020 (Linnean Society of London and the Systematics Association). RF acknowledges support from the following sources of funding: Ramón y Cajal fellowship (grant agreement no. RYC-2017-22492 funded by MCIN/AEI/10.13039/501100011033 and ESF ‘Investing in your future’), project PID2019-108824GA-I00 funded by MCIN/AEI/10.13039/501100011033, and by the European Research Council (ERC) under the European’s Union’s Horizon 2020 research and innovation programme (grant agreement no. 948281)). SJBC acknowledges support from the Australian Research Council (grant agreements: DP230100731, DP18010385 and DP120102132). We acknowledge Ignacio Ribera for his invaluable contribution at the beginning of this project and for supervising this research until the end of his life. We also thank Centro de Supercomputación de Galicia (CESGA) for access to computer resources, and acknowledge the anonymous reviewers for their suggestions.

## 7. CONTRIBUTIONS

PBG and RF designed and conceived the research. PBG, SP and SJBC provided valuable knowledge about the studied systems. TMB and PGBH generated transcriptomic data of the aquatic lineages. PBG generated genomic data of the terrestrial lineages, processed both newly generated and public genomic data and performed all analyses. PBG prepared the figures and other provided materials. PBG, SJBC and RF interpreted and discussed the results. SJBC and RF provided resources. RF supervised the study. PBG and RF drafted the paper. All authors contributed to the final version of the manuscript.

## SUPPLEMENTARY FIGURES

**Supplementary Figure 1.**
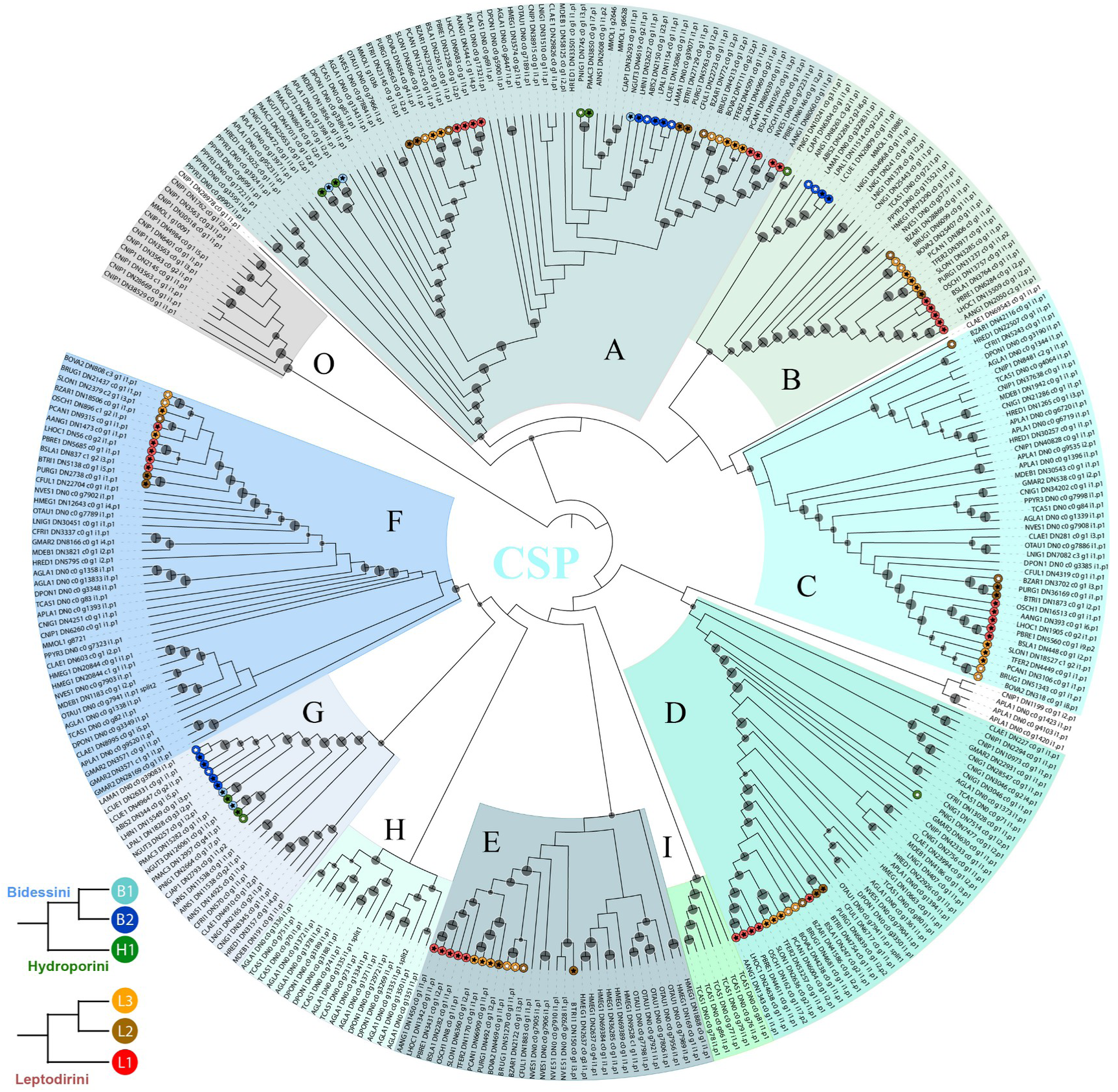
Maximum-likelihood phylogenetic tree for chemosensory proteins (CSP). Colored ranges indicate significantly supported clades (i.e. bootstrap >75%). “O” clade corresponds to a clade composed exclusively by non-coleopteran CSPs (i.e., the Strepsiptera species *Mengenilla moldrzyki* (MMOL1) and the Neuroptera species *Chrysoperla nipponensis* (CNIP1)). “A–I” correspond to supported clades also including coleopteran CSPs.

**Supplementary Figure 2.**
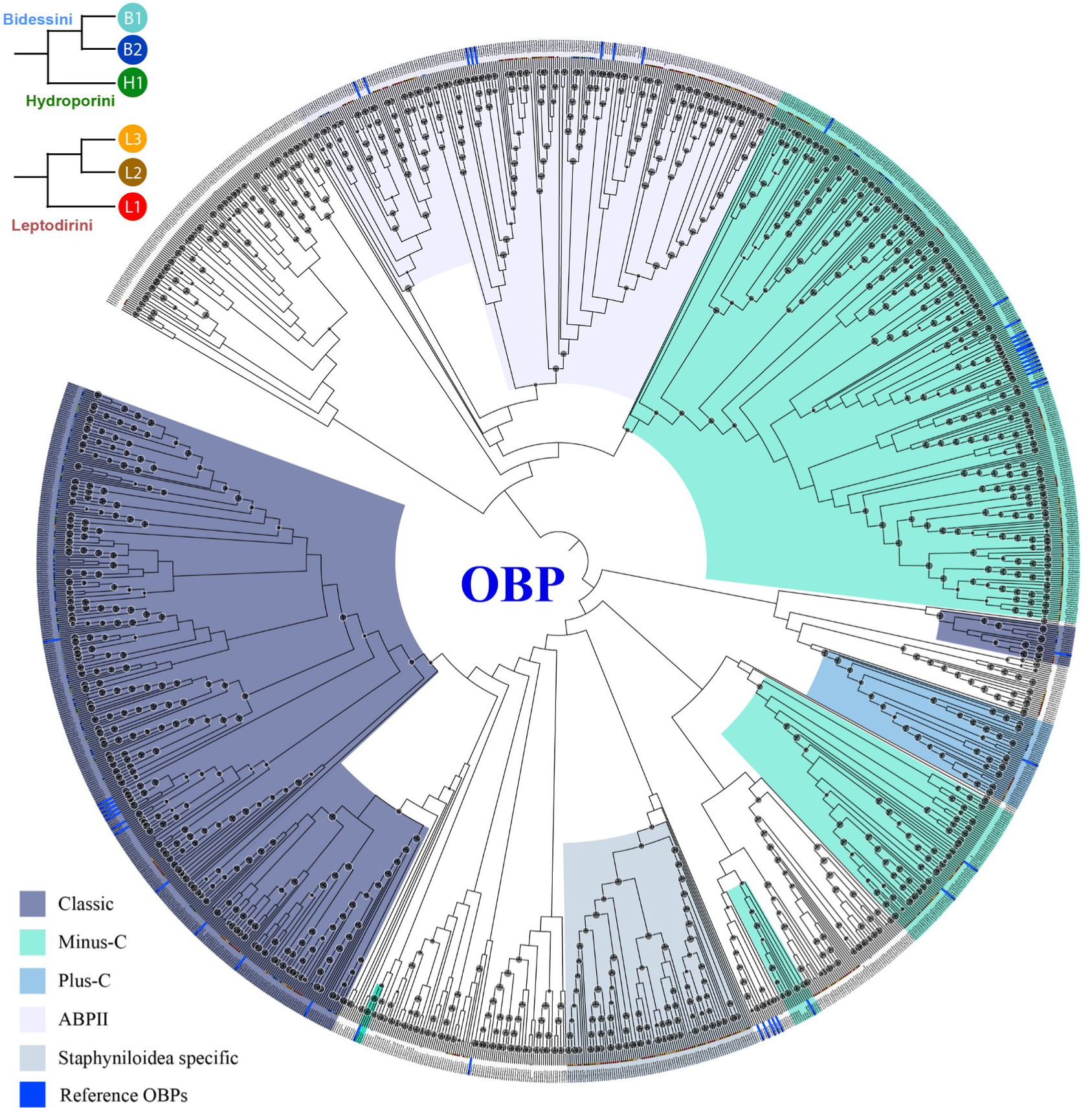
Maximum-likelihood phylogenetic tree for odorant binding proteins (OBP). Colored ranges indicate significantly supported clades (i.e. bootstrap >75%). Dark blue tagged tips correspond to OBPs previously described in *Tribolium castaneum* (Tcas) used as a guide to identify the main OBP clades in Coleoptera (e.g., Minus-C OBPs, Plus-C OBPs, antennal binding protein II (ABPII); Dippel et al. 2014).

**Supplementary Figure 3.**
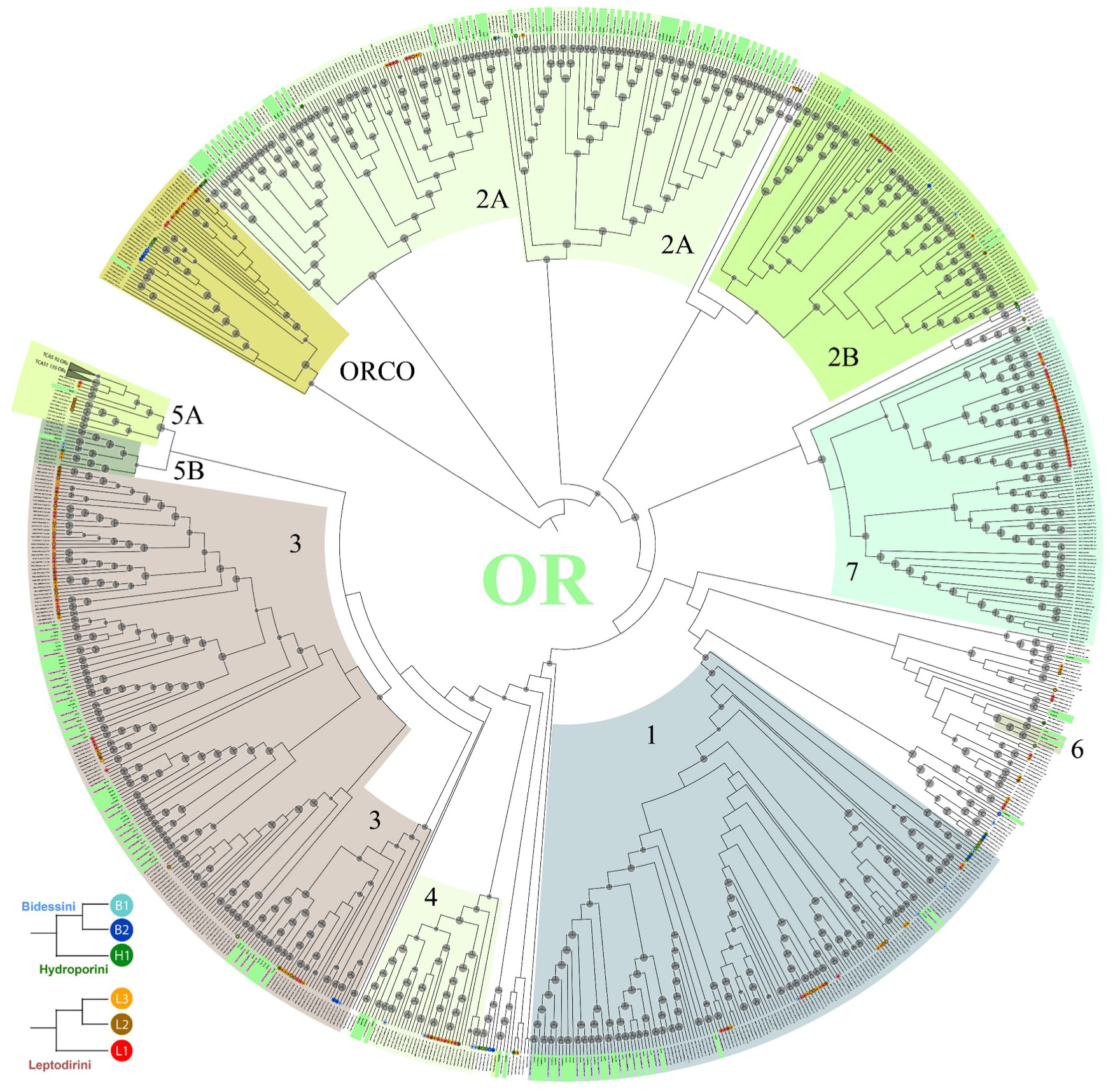
Maximum-likelihood phylogenetic tree for odorant receptors (OR). Colored ranges indicate significantly supported clades (i.e. bootstrap >75%). Green tagged tips correspond to previously described ORs in *Tribolium castaneum* (Tcas) (Dippel et al. 2016) used as a guide to identify the main OR clades in Coleoptera (Andersson et al. 2019). The lineage-specific OR5A clades of *Tribolium castaneum* are collapsed (i.e., 227 ORs).

**Supplementary Figure 4.**
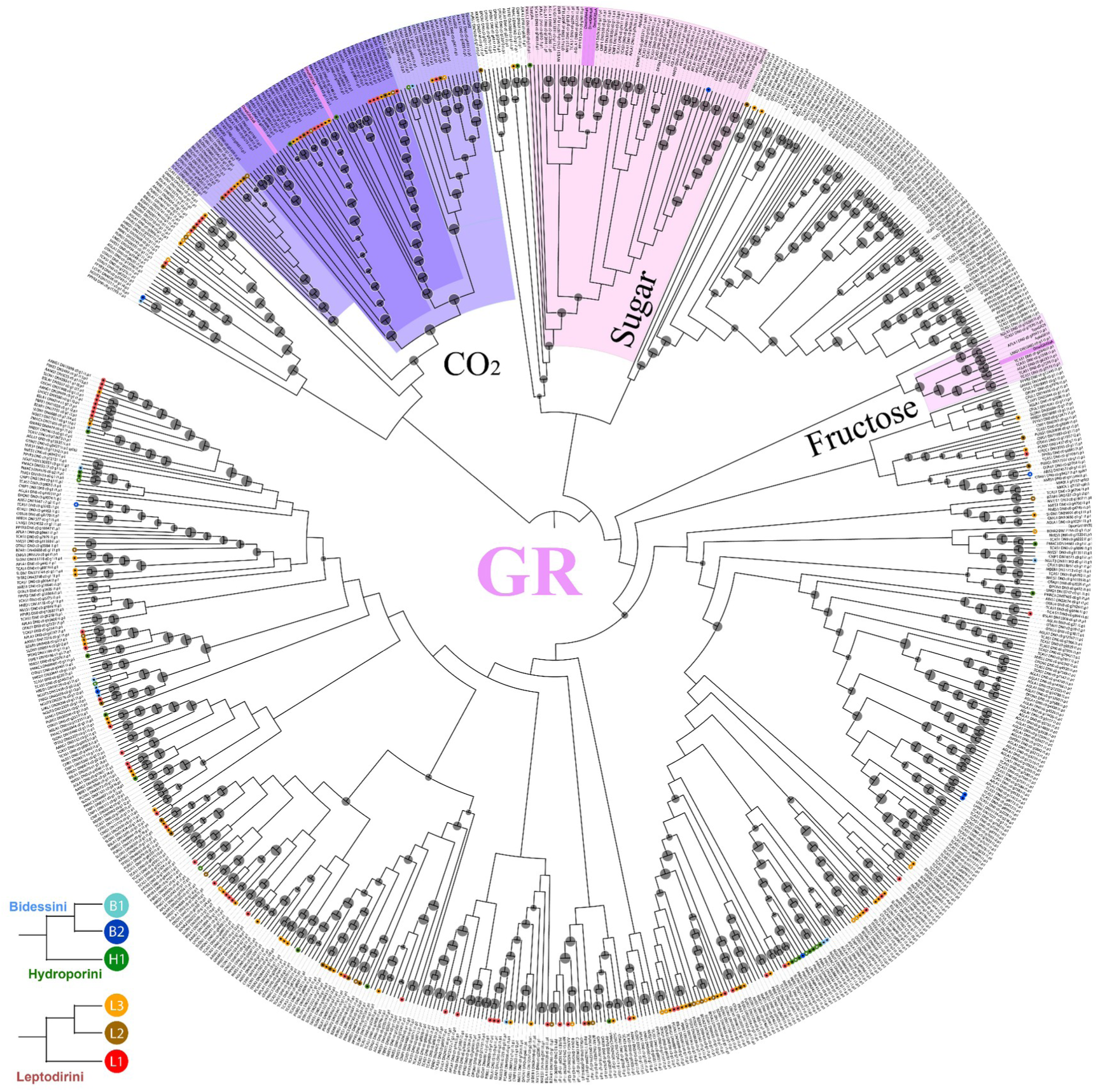
Maximum-likelihood phylogenetic tree for gustatory receptors (GR). Colored ranges indicate significantly supported clades (i.e. bootstrap >75%). Pink tagged tips correspond to GRs previously described in *Drosophila melanogaster* (Dmel) used as a guide to identify the main GR clades in insects (flybase.org). Previously described GRs in *Tribolium castaneum* (Tcas) (Dippel et al. 2016), *Agrilus planipennis* (Apla) and *Dendroctonus ponderosae* (Dpon) (Andersson et al. 2019) were also added as a reference to detect the conserved GR clades in Coleoptera. The two previously described clades of CO_2_ receptors are colored in dark purple, including the *D. melanogaster* GR21 and GR63. A third CO_2_ receptor clade and a sister clade to all these the CO_2_ receptors are colored in light purple.

**Supplementary Figure 5.**
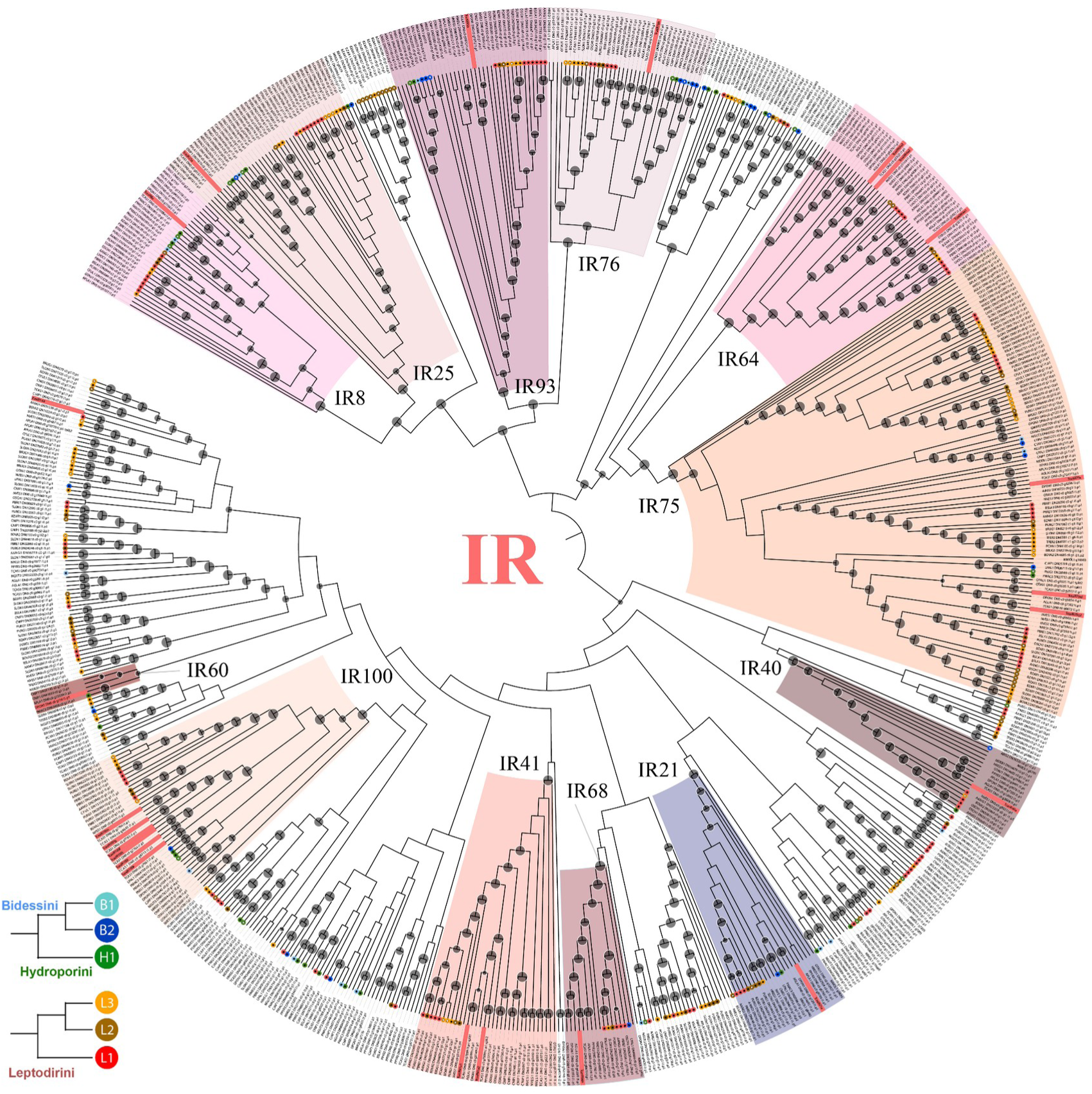
Maximum-likelihood phylogenetic tree for ionotropic receptors (IR). Colored ranges indicate significantly supported clades (i.e. bootstrap >75%). Red tagged tips correspond to IRs previously described in *Tribolium castaneum* (Tcas) (Dippel et al. 2016) used as a guide to identify the main IR clades in Coleoptera (Andersson et al. 2019).

**Supplementary Figure 6.**
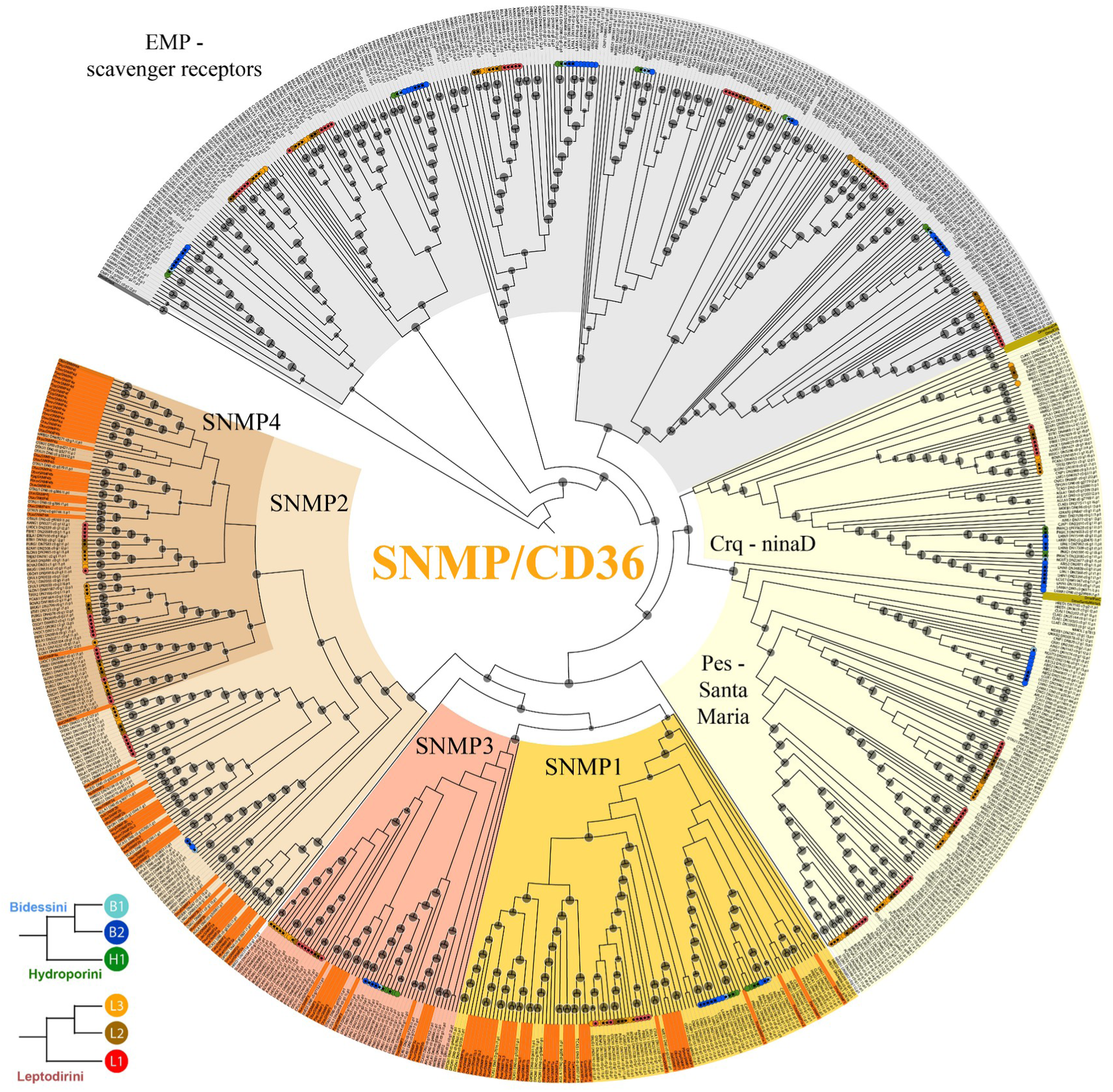
Maximum-likelihood phylogenetic tree for sensory neuron membrane proteins (SNMP) and other conserved clades of the Cluster of Differentiation 36 (CD36) protein superfamily (i.e., EMP-scavenger receptors, Crq, ninaD, Pes, Santa Maria). Colored ranges indicate significantly supported clades (i.e. bootstrap >75%). Grey and gold tagged tips correspond to previously described CD36 proteins in *Drosophila melanogaster* (Dmel) (flybase.org). Orange tagged tips correspond to previously described SNMPs in *D. melanogaster* and in several Coleoptera species (e.g. *Aethina tumida* (Atum), *Nicrophorus vespilloides* (Nves), *Onthophagus taurus* (Otau), *Photinus pyralis* (Ppyr), *Rhaphuma horsfieldi* Rhor) obtained from Zhao et al. (2020). These previously described SNMP/CD36 sequences were used as a guide to identify supported clades in Coleoptera.

**Supplementary Figure 7.**
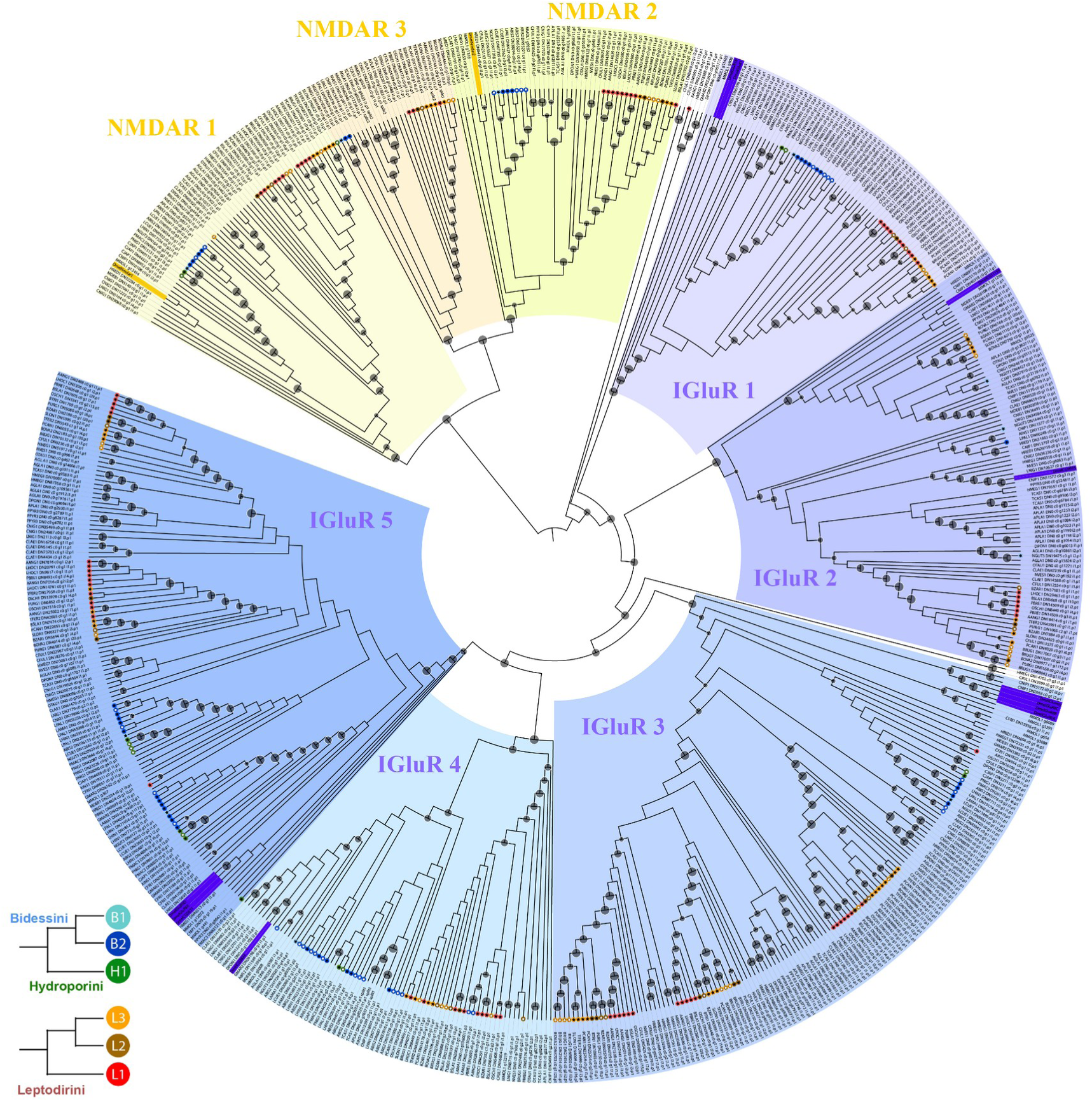
Maximum-likelihood phylogenetic tree for ionotropic glutamate receptors (IGluR). Colored ranges indicate significantly supported clades (i.e. bootstrap >75%). Yellow and purple tagged tips correspond to previously described N-methyl-D-aspartate receptors (NMDARs) and IGluRs in *Drosophila melanogaster* (Dmel) respectively, used as a guide to identify these clades in Coleoptera. IGluR1–5 correspond to significantly supported clades of IGluRs in our study.

## SUPPLEMENTARY TABLE CAPTIONS

**Supplementary Table 1.** Chemosensory gene count annotated in each species for each chemosensory gene family, including the following information: data type (genome/transcriptome), ecology (aquatic/terrestrial), habitat type (surface/subterranean) and lineage (Outgroup, Archostemata, Myxophaga, Adephaga, Polyphaga).

**Supplementary Table 2.** BadiRate results for the following models: global rates, ancestral rates, habitat shift rates, subterranean branch rates, highly modified lineages rates. Results are indicated for the all taxa analysis, the Leptodirini analysis and the Hydroporini-Bidessini analysis.

## REFERENCES

1. Accordi, F.S., Sbordoni, V., 1977. The fine structure of Hamann’s organ in *Leptodirus hochenwartii*, a highly specialized cave bathysciinae: (Coleoptera, Catopidae). Int. J. Speleol. 9:153.165.

2. Almudi, I., Vizueta, J., Wyatt, C.D.R., de Mendoza, A., Marlétaz, F., Firbas, P.N., Feuda, R., Masiero, G., Medina, P., Alcaina-Caro, A., Cruz, F., Gómez-Garrido, J., Gut, M., Alioto, T.S., Vargas-Chavez, C., Davie, K., Misof, B., González, J., Aerts, S., Lister, R., Paps, J., Rozas, J., Sánchez-Gracia, A., Irimia, M., Maeso, I., Casares, F., 2020. Genomic adaptations to aquatic and aerial life in mayflies and the origin of insect wings. Nat. Commun. 11, 2631.

3. Andersson, M.N., Keeling, C.I., Mitchell, R.F., 2019. Genomic content of chemosensory genes correlates with host range in wood-boring beetles (*Dendroctonus ponderosae*, Agrilus planipennis, and Anoplophora glabripennis). BMC Genomics. 20.

4. Balart-García, P., Cieslak, A., Escuer, P., Rozas, J., Ribera, I., Fernández, R., 2021. Smelling in the dark: Phylogenomic insights into the chemosensory system of a subterranean beetle. Mol. Ecol. 30, 2573–2590.

5. Balart-García, P., Aristide, L., Bradford, T.M., Beasley-Hall, P.G., Polak, S., Cooper, S.J.B., Fernández, R., 2023. Parallel and convergent genomic changes underlie independent subterranean colonization across beetles. Nat. Commun. 14, 3842.

6. Benton, R., 2015. Multigene Family Evolution: Perspectives from Insect Chemoreceptors. Trends Ecol. Evol. 30, 590–600.

7. Benton, R., Vannice, K.S., Gomez-Diaz, C., Vosshall, L.B., 2009. Variant ionotropic glutamate receptors as chemosensory receptors in *Drosophila*. Cell 136, 149–162.

8. Blum, M., Chang, H.-Y., Chuguransky, S., Grego, T., Kandasaamy, S., Mitchell, A., Nuka, G., Paysan-Lafosse, T., Qureshi, M., Raj, S., Richardson, L., Salazar, G.A., Williams, L., Bork, P., Bridge, A., Gough, J., Haft, D.H., Letunic, I., Marchler-Bauer, A., Mi, H., Natale, D.A., Necci, M., Orengo, C.A., Pandurangan, A.P., Rivoire, C., Sigrist, C.J.A., Sillitoe, I., Thanki, N., Thomas, P.D., Tosatto, S.C.E., Wu, C.H., Bateman, A., Finn, R.D., 2021. The InterPro protein families and domains database: 20 years on. Nucleic Acids Res. 49, D344–D354.

9. Buchfink, B., Reuter, K., Drost, H.-G., 2021. Sensitive protein alignments at tree-of-life scale using DIAMOND. Nat. Methods 18, 366–368.

10. Capella-Gutiérrez, S., Silla-Martínez, J.M., Gabaldón, T., 2009. trimAl: a tool for automated alignment trimming in large-scale phylogenetic analyses. Bioinformatics 25, 1972–1973.

11. Challis, R., Richards, E., Rajan, J., Cochrane, G., Blaxter, M., 2020. BlobToolKit – Interactive Quality Assessment of Genome Assemblies. G3 10, 1361–1374.

12. Chiang, A.-S., Lin, W.-Y., Liu, H.-P., Pszczolkowski, M.A., Fu, T.-F., Chiu, S.-L., Holbrook, G.L., 2002. Insect NMDA receptors mediate juvenile hormone biosynthesis. Proc. Natl. Acad. Sci. U. S. A. 99, 37–42.

13. Cieslak, A., Fresneda, J., Ribera, I., 2014. Life-history specialization was not an evolutionary dead-end in Pyrenean cave beetles. Proceedings of the Royal Society B: Biological Sciences. 281:20132978.

14. Clyne, P.J., Warr, C.G., Freeman, M.R., Lessing, D., Kim, J., Carlson, J.R., 1999. A novel family of divergent seven-transmembrane proteins: candidate odorant receptors in *Drosophila*. Neuron 22, 327–338.

15. Cohen, A.C., Cheah, C.A., 2015. Interim diets for specialist predators of hemlock woolly adelgids. Entomology, Ornithology & Herpetology 4, 1.

16. Cooper, S.J.B., Hinze, S., Leys, R., Watts, C.H.S., Humphreys, W.F., 2002. Islands under the desert: molecular systematics and evolutionary origins of stygobitic water beetles (Coleoptera: Dytiscidae) from central Western Australia. Invertebrate Systematics 16, 589–598.

17. Delay, B., 1978. Milieu souterrain et écophysiologie de la reproduction et du développement des Coléoptères Bathysciinae hypogés. Mém Biospéol 5, 1–349.

18. Deleurance-Glaçon, S., 1963. Recherches sur les Coléoptères troglobies de la sous-famille Bathysciinae. Annu Sci Nat Zool Paris 12, 1–172.

19. Dippel, S., Oberhofer, G., Kahnt, J., Gerischer, L., Opitz, L., Schachtner, J., Stanke, M., Schütz, S., Wimmer, E. A., Angeli, S., 2014. Tissue-specific transcriptomics, chromosomal localization, and phylogeny of chemosensory and odorant binding proteins from the red flour beetle *Tribolium castaneum* reveal subgroup specificities for olfaction or more general functions. BMC Genomics 15:1141

20. Dippel, S., Kollmann, M., Oberhofer, G., Montino, A., Knoll, C., Krala, M., Rexer, K.-H., Frank, S., Kumpf, R., Schachtner, J., Wimmer, E.A., 2016. Morphological and Transcriptomic Analysis of a Beetle Chemosensory System Reveals a Gnathal Olfactory Center. BMC Biol. 14, 90.

21. Eyun, S.-I., Soh, H.Y., Posavi, M., Munro, J.B., Hughes, D.S.T., Murali, S.C., Qu, J., Dugan, S., Lee, S.L., Chao, H., Dinh, H., Han, Y., Doddapaneni, H., Worley, K.C., Muzny, D.M., Park, E.-O., Silva, J.C., Gibbs, R.A., Richards, S., Lee, C.E., 2017. Evolutionary History of Chemosensory-Related Gene Families across the Arthropoda. Mol. Biol. Evol. 34, 1838– 1862.

22. Fernández, R., Tonzo, V., Guerrero, C.S., Lozano-Fernandez, J., Martínez-Redondo, G.I., Balart-García, P., Aristide, L., Eleftheriadi, K., Vargas-Chávez, C., 2022. MATEdb, a data repository of high-quality metazoan transcriptome assemblies to accelerate phylogenomic studies. Peer Community Journal 2.

23. Fresneda, J., Grebennikov, V.V., Ribera, I., 2011. The phylogenetic and geographic limits of Leptodirini (Insecta: Coleoptera: Leiodidae: Cholevinae), with a description of Sciaphyes shestakovi sp. n. from the Russian Far East. Arthropod Syst. Phylogeny. 69.

24. Fu, L., Niu, B., Zhu, Z., Wu, S., Li, W., 2012. CD-HIT: accelerated for clustering the next-generation sequencing data. Bioinformatics 28, 3150–3152.

25. Gibert, J., Deharveng, L., 2002. Subterranean ecosystems: a truncated functional biodiversity. BioScience 52:473–481.

26. Gomez-Diaz, C., Bargeton, B., Abuin, L., Bukar, N., Reina, J.H., Bartoi, T., Graf, M., Ong, H., Ulbrich, M.H., Masson, J.-F., Benton, R., 2016. A CD36 ectodomain mediates insect pheromone detection via a putative tunnelling mechanism. Nat. Commun. 7, 11866.

27. Hoang, D.T., Chernomor, O., von Haeseler, A., Minh, B.Q., Vinh, L.S., 2018. UFBoot2: Improving the Ultrafast Bootstrap Approximation. Mol. Biol. Evol. 35, 518–522.

28. Hüppop, K., 2012. Adaptation to low food. In: White WB, Culver DC, editors. Encyclopedia of caves. Academic Press, Waltham, MA, p. 1–9.

29. Joseph, R.M., Carlson, J.R., 2015. Drosophila Chemoreceptors: A Molecular Interface Between the Chemical World and the Brain. Trends Genet. 31, 683–695.

30. Kane, T.C., Poulson, T.L., 1976. Foraging by Cave Beetles: Spatial and Temporal Heterogeneity of Prey. Ecology. 57:793–800.

31. Katoh, K., Standley, D.M., 2013. MAFFT multiple sequence alignment software version 7: improvements in performance and usability. Mol. Biol. Evol. 30, 772–780.

32. Knecht, Z.A., Silbering, A.F., Cruz, J., Yang, L., Croset, V., Benton, R., Garrity, P.A., 2017. Ionotropic Receptor-dependent moist and dry cells control hygrosensation in. Elife 6.

33. Kováč, Ľ., 2018. Caves as Oligotrophic Ecosystems, in: Moldovan, O.T., Kováč, Ľ., Halse, S. (Eds.), Cave Ecology. Springer International Publishing, Cham, pp. 297–307.

34. Kozma, M.T., Ngo-Vu, H., Wong, Y.Y., Shukla, N.S., Pawar, S.D., Senatore, A., Schmidt, M., Derby, C.D., 2020. Comparison of transcriptomes from two chemosensory organs in four decapod crustaceans reveals hundreds of candidate chemoreceptor proteins. PLoS One 15, e0230266.

35. Langille, B.L., Hyde, J., Saint, K.M., Bradford, T.M., Stringer, D.N., Tierney, S.M., Humphreys, W.F., Austin, A.D., Cooper, S.J.B., 2021. Evidence for speciation underground in diving beetles (Dytiscidae) from a subterranean archipelago. Evolution 75, 166–175.

36. Lauritzen, S.-E., 2018. Physiography of the Caves, in: Moldovan, O.T., Kováč, Ľ., Halse, S. (Eds.), Cave Ecology. Springer International Publishing, Cham, pp. 7–21.

37. Lawrence, J.F., 2016. 10.4. Scirtidae Fleming, 1821. In: Beutel, R.G., Leschen, R.A.B. (eds) Handbook of zoology, vol 4/38: Coleoptera, beetles, vol 1. Morphology and systematics (Archostemata, Adephaga, Myxophaga, Polyphaga partim), 2nd edn. W DeGruyter, Berlin, pp 215–225

38. Leijs, R., van Nes, E.H., Watts, C.H., Cooper, S.J.B., Humphreys, W.F., Hogendoorn, K., 2012. Evolution of blind beetles in isolated aquifers: a test of alternative modes of speciation. PLoS One 7, e34260.

39. Letunic, I., Bork, P., 2021. Interactive Tree Of Life (iTOL) v5: an online tool for phylogenetic tree display and annotation. Nucleic Acids Res. 49, W293–W296.

40. Leys, R., Watts, C.H.S., Cooper, S.J.B., Humphreys, W.F., 2003. Evolution of subterranean diving beetles (Coleoptera: Dytiscidae: Hydroporini, Bidessini) in the arid zone of Australia. Evolution 57, 2819–2834.

41. Librado, P., Vieira, F.G., Rozas, J., 2012. BadiRate: estimating family turnover rates by likelihood-based methods. Bioinformatics 28:279–281.

42. Manni, M., Berkeley, M.R., Seppey, M., Simão, F.A., Zdobnov, E.M., 2021. BUSCO Update: Novel and Streamlined Workflows along with Broader and Deeper Phylogenetic Coverage for Scoring of Eukaryotic, Prokaryotic, and Viral Genomes. Mol. Biol. Evol. 38:4647–4654.

43. Mayer, M.L., Armstrong, N., 2004. Structure and Function of Glutamate Receptor Ion Channels. Annual Review of Physiology 66:161–181.

44. McKenna, D.D., Shin, S., Ahrens, D., Balke, M., Beza-Beza, C., Clarke, D.J., Donath, A., Escalona, H.E., Friedrich, F., Letsch, H., Liu, S., Maddison, D., Mayer, C., Misof, B., Murin, P.J., Niehuis, O., Peters, R.S., Podsiadlowski, L., Pohl, H., Scully, E.D., Yan, E.V., Zhou, X., Ślipiński, A., Beutel, R.G., 2019. The evolution and genomic basis of beetle diversity. Proc. Natl. Acad. Sci. U. S. A. 116, 24729–24737.

45. Minh, B.Q., Schmidt, H.A., Chernomor, O., Schrempf, D., Woodhams, M.D., von Haeseler, A., Lanfear, R., 2020. IQ-TREE 2: New Models and Efficient Methods for Phylogenetic Inference in the Genomic Era. Mol. Biol. Evol. 37, 1530–1534.

46. Mitchell, R.F., & Andersson, M.N., 2021. 17 – Olfactory genomics of the Coleoptera. In G. J. Blomquist & R. G. Vogt (Eds.), Insect Pheromone Biochemistry and Molecular Biology (Second Edition) (pp. 547–590). Academic Press.

47. Mitchell, R.F., Schneider, T.M., Schwartz, A.M., Andersson, M.N., McKenna, D.D., 2020. The diversity and evolution of odorant receptors in beetles (Coleoptera). Insect Mol. Biol. 29: 77–91.

48. Moldovan, O.T., Jalžić, B., Erichsen, E., 2004. Adaptation of the mouthparts in some subterranean Cholevinae (Coleoptera, Leiodidae). Natura Croatica 13:1–18.

49. Ngoc, P.C.T., Greenhalgh, R., Dermauw, W., Rombauts, S., Bajda, S., Zhurov, V., Grbić, M., Van de Peer, Y., Van Leeuwen, T., Rouzé, P., Clark, R.M., 2016. Complex Evolutionary Dynamics of Massively Expanded Chemosensory Receptor Families in an Extreme Generalist Chelicerate Herbivore. Genome Biol. Evol. 8, 3323–3339.

50. Nichols, Z., Vogt, R.G., 2008. The SNMP/CD36 gene family in Diptera, Hymenoptera and Coleoptera: *Drosophila melanogaster*, *D. pseudoobscura*, *Anopheles gambiae*, *Aedes aegypti, Apis mellifera, and Tribolium castaneum*. Insect Biochem. Mol. Biol. 38, 398–415.

51. Ni, L., 2020. The Structure and Function of Ionotropic Receptors in *Drosophila*. Front. Mol. Neurosci. 13, 638839.

52. Peck, S. B., 1984. The distribution and evolution of cavernicolous *Ptomaphagus* beetles in the southeastern United States (Coleoptera; Leiodidae; Cholevinae) with new species and records. Canadian Journal of Zoology, 62(4): 730–740.

53. Pelosi, P., Iovinella, I., Felicioli, A., Dani, F.R., 2014. Soluble proteins of chemical communication: an overview across arthropods. Front. Physiol. 5, 320.

54. Pelosi, P., Iovinella, I., Zhu, J., Wang, G., Dani, F.R., 2018. Beyond chemoreception: diverse tasks of soluble olfactory proteins in insects. Biological Reviews 93:184–200.

55. Poulson, T.L., 1963. Cave Adaptation in Amblyopsid Fishes. American Midland Naturalist 70:257.

56. Price, M.N., Dehal, P.S., Arkin, A.P., 2010. FastTree 2--approximately maximum-likelihood trees for large alignments. PLoS One 5, e9490.

57. Ribera, I., Fresneda, J., Bucur, R., Izquierdo, A., Vogler, A.P., Salgado, J.M., Cieslak, A., 2010. Ancient origin of a Western Mediterranean radiation of subterranean beetles. BMC Evol. Biol. 10, 29.

58. Robertson, H.M., Baits, R.L., Walden, K.K.O., Wada-Katsumata, A., Schal, C., 2018. Enormous expansion of the chemosensory gene repertoire in the omnivorous German cockroach *Blattella germanica*. J. Exp. Zool. B Mol. Dev. Evol. 330, 265–278.

59. Robertson, H.M., Warr, C.G., Carlson, J.R., 2003. Molecular evolution of the insect chemoreceptor gene superfamily in *Drosophila melanogaster*. Proc. Natl. Acad. Sci. U. S. A. 100:14537–14542.

60. Sambrook, J., Fritsch, E.F., Maniatis, T., 1989. Molecular Cloning: A Laboratory Manual. Cold Spring Harbor Laboratory Press.

61. Sánchez-Gracia, A., Vieira, F.G., Rozas, J., 2009. Molecular evolution of the major chemosensory gene families in insects. Heredity 103:208–216.

62. Schoville, S.D., Chen, Y.H., Andersson, M.N., Benoit, J.B., Bhandari, A., Bowsher, J.H., Brevik, K., Cappelle, K., Chen, M.-J.M., Childers, A.K., Childers, C., Christiaens, O., Clements, J., Didion, E.M., Elpidina, E.N., Engsontia, P., Friedrich, M., García-Robles, I., Gibbs, R.A., Goswami, C., Grapputo, A., Gruden, K., Grynberg, M., Henrissat, B., Jennings, E.C., Jones, J.W., Kalsi, M., Khan, S.A., Kumar, A., Li, F., Lombard, V., Ma, X., Martynov, A., Miller, N.J., Mitchell, R.F., Munoz-Torres, M., Muszewska, A., Oppert, B., Palli, S.R., Panfilio, K.A., Pauchet, Y., Perkin, L.C., Petek, M., Poelchau, M.F., Record, É., Rinehart, J.P., Robertson, H.M., Rosendale, A.J., Ruiz-Arroyo, V.M., Smagghe, G., Szendrei, Z., Thomas, G.W.C., Torson, A.S., Vargas Jentzsch, I.M., Weirauch, M.T., Yates, A.D., Yocum, G.D., Yoon, J.-S., Richards, S., 2018. A model species for agricultural pest genomics: the genome of the Colorado potato beetle, *Leptinotarsa decemlineata* (Coleoptera: Chrysomelidae). Sci. Rep. 8, 1931.

63. Stegner, M.E.J., Stemme, T., Iliffe, T.M., Richter, S., Wirkner, C.S., 2015. The brain in three crustaceans from cavernous darkness. BMC Neurosci. 16, 19.

64. Stocker, R.F., 1994. The organization of the chemosensory system in *Drosophila melanogaster*: a review. Cell Tissue Res. 275, 3–26.

65. Thoma, M., Missbach, C., Jordan, M.D., Grosse-Wilde, E., Newcomb, R.D., Hansson, B.S., 2019. Transcriptome Surveys in Silverfish Suggest a Multistep Origin of the Insect Odorant Receptor Gene Family. Front. Ecol. Evol. 7

66. van Giesen, L., Garrity, P.A., 2017. More than meets the IR: the expanding roles of variant Ionotropic Glutamate Receptors in sensing odor, taste, temperature and moisture. F1000Res. 6, 1753.

67. Varatharasan, N., Croll, R.P., Franz-Odendaal, T., 2009. Taste bud development and patterning in sighted and blind morphs of *Astyanax mexicanus*. Dev. Dyn. 238, 3056–3064.

68. Vizueta, J., Frías-López, C., Macías-Hernández, N., Arnedo, M.A., Sánchez-Gracia, A., Rozas, J., 2017. Evolution of Chemosensory Gene Families in Arthropods: Insight from the First Inclusive Comparative Transcriptome Analysis across Spider Appendages. Genome Biol. Evol. 9, 178–196.

69. Vizueta, J., Sánchez-Gracia, A., Rozas, J., 2020. bitacora: A comprehensive tool for the identification and annotation of gene families in genome assemblies. Mol. Ecol. Resour. 20, 1445–1452.

70. Wang, H.-C., Minh, B.Q., Susko, E., Roger, A.J., 2018. Modeling Site Heterogeneity with Posterior Mean Site Frequency Profiles Accelerates Accurate Phylogenomic Estimation. Systematic Biology 67:216–235.

71. Watts C.H.S & Leys R., 2008. Review of the epigean species of Paroster SHARP, 1882, with descriptions of three new species, and phylogeny based on DNA sequence data of two mitochondrial genes (Coleoptera: Dytiscidae: Hydroporinae). Koleopterol. Rundsch. 78, 9–36.

72. Yang, J., Chen, X., Bai, J., Fang, D., Qiu, Y., Jiang, W., Yuan, H., Bian, C., Lu, J., He, S., Pan, X., Zhang, Y., Wang, X., You, X., Wang, Y., Sun, Y., Mao, D., Liu, Y., Fan, G., Zhang, H., Chen, X., Zhang, X., Zheng, L., Wang, J., Cheng, L., Chen, J., Ruan, Z., Li, J., Yu, H., Peng, C., Ma, X., Xu, J., He, Y., Xu, Z., Xu, P., Wang, J., Yang, H., Wang, J., Whitten, T., Xu, X., Shi, Q., 2016. The *Sinocyclocheilus* cavefish genome provides insights into cave adaptation. BMC Biol. 14, 1.

73. Yoshizawa, M., Jeffery, W.R., van Netten, S.M., McHenry, M.J., 2014. The sensitivity of lateral line receptors and their role in the behavior of Mexican blind cavefish (*Astyanax mexicanus*). J. Exp. Biol. 217, 886–895.

74. Zhao, Y., Guzik M.T., Humphreys, W.F., Watts, C.H.S., Cooper, S.J.B., Sherratt, E., 2023. Evolutionary transition from surface to subterranean living in Australian water beetles (Coleoptera, Dytiscidae) through adaptive and relaxed selection. Biol. J. Linn. Soc. 10.1093/biolinnean/blad142.

75. Zhao, Y.-J., Li, G.-C., Zhu, J.-Y., Liu, N.-Y., 2020. Genome-based analysis reveals a novel SNMP group of the Coleoptera and chemosensory receptors in *Rhaphuma horsfieldi*. Genomics 112:2713–2728.

## SUPPLEMENTARY REFERENCES

1. Andersson, M.N., Keeling, C.I., Mitchell, R.F., 2019. Genomic content of chemosensory genes correlates with host range in wood-boring beetles (*Dendroctonus ponderosae*, Agrilus planipennis, and Anoplophora glabripennis). BMC Genomics. 20.

2. Dippel, S., Oberhofer, G., Kahnt, J., Gerischer, L., Opitz, L., Schachtner, J., Stanke, M., Schütz, S., Wimmer, E. A., Angeli, S., 2014. Tissue-specific transcriptomics, chromosomal localization, and phylogeny of chemosensory and odorant binding proteins from the red flour beetle *Tribolium castaneum* reveal subgroup specificities for olfaction or more general functions. BMC Genomics 15:1141

3. Dippel, S., Kollmann, M., Oberhofer, G., Montino, A., Knoll, C., Krala, M., Rexer, K.-H., Frank, S., Kumpf, R., Schachtner, J., Wimmer, E.A., 2016. Morphological and Transcriptomic Analysis of a Beetle Chemosensory System Reveals a Gnathal Olfactory Center. BMC Biol. 14, 90.

4. Zhao, Y.-J., Li, G.-C., Zhu, J.-Y., Liu, N.-Y., 2020. Genome-based analysis reveals a novel SNMP group of the Coleoptera and chemosensory receptors in *Rhaphuma horsfieldi*. Genomics 112:2713–2728.

